# Disentangling the temporal signatures of conditioned pupil dilation: Distinct valence- and prediction-error-related components revealed by mega-analysis

**DOI:** 10.64898/2026.07.16.737017

**Authors:** Johannes B. Finke, Anna M. Schippers, Hartmut Schächinger, Tim Klucken

## Abstract

Pupil dilation offers a sensitive, non-invasive window into the neurocognitive processes supporting human associative learning. Yet, the field lacks consensus on how to quantify conditioned pupil responses, particularly which time intervals best serve as readouts. Moreover, it remains unresolved whether pupil dilation during learning reflects a unitary arousal signal or multiple (overlapping yet distinct) processes that unfold sequentially (e.g., cue valuation, uncertainty-driven attention, anticipation, prediction-error [PE] signaling).

To address this, we conducted a large-scale individual-participant-data reanalysis (mega-analysis) of harmonized primary studies from our lab (*K* = 7 independent samples; total *N* = 385; 562 sessions), reprocessed through an identical pipeline, with systematic variation in outcome valence and modality. Temporal PCA of epochs locked either to the conditioned (CS) or unconditioned stimulus (US) yielded highly stable component structures across trials and studies, which carried dissociable information: An early (CS-locked) component selectively tracked appetitive value, distinguishing reward cues before aversive differentiation emerged. Later phases reflected differential conditioning across both valences, increasingly coupled to arousal toward US onset. Aversive learning was most specifically expressed in responses peaking around expected US delivery, which also predicted negative valence ratings. Two subsequent components captured unconditioned responding and its modulation by unexpected outcomes and omissions, consistent with unsigned PE signaling (amplified after aversive cues) and, later, positive PEs. Generalized additive mixed models largely converged with the PCA structure.

Pupil dilation is therefore not a unitary marker of conditioning, but rather a high-resolution readout of sequential processes such as valuation, arousal/attention, and outcome updating, offering a promising framework for future computational models of learning.

## 1. Introduction

As a non-invasive index of central neurocognitive processes, pupil dilation has been highlighted as a promising marker of differential Pavlovian conditioning in humans (Finke et al., 2021), with the potential to track associative learning over time (Pietrock et al., 2019). Yet, approaches to quantifying conditioned pupil dilation vary widely between studies, especially in the time windows extracted for analysis, which may blur temporal information inherent in the signal, and thus limit the inferences that can be drawn from it. Moreover, while noradrenergic arousal (Aston-Jones & Cohen, 2005; Joshi et al., 2016) has been proposed as a general underlying mechanism (but see Megemont et al., 2022), and there is evidence for pupil-size modulation by surprise and prediction-error processing in a reinforcement-learning context (Preuschoff et al., 2011; de Berker et al., 2016; Van Slooten et al., 2018), very little is known about the specific functional significance of pupil dilation during Pavlovian conditioning, and whether this also depends on latent (within-trial) temporal dynamics. Several candidate processes have been formalized in computational models of associative learning (e.g., Rescorla-Wagner [RW], Pearce-Hall [PH]; Rescorla & Wagner, 1972; Pearce & Hall, 1980), which distinguish learning quantities such as associative strength, signed or unsigned prediction error, and associability/attentional weight. There is initial evidence that some of these model-derived signals covary with pupil dilation during Pavlovian learning (Pietrock et al., 2019; Finke et al., 2023b), yet it is unclear whether they also map onto separable components of the pupil response.

Recent studies have indeed begun to link pupil-size modulation at different latencies to distinct types of learning signals, such as salience, feedback processing, and memory updating (Van Slooten et al., 2018; Colizoli et al., 2026), raising the possibility that response latency itself carries functional information about underlying computations, but findings from aversive/fear conditioning studies remain inconsistent (Stemerding et al., 2022; Liu et al., 2026), and between-study heterogeneity in methodological choices is likely to contribute to this lack of understanding. Beyond value-sensitive responses to reward-predictive cues (Schad et al., 2020), pupil dilation during Pavlovian learning has been suggested to index either outcome expectancy (Tzovara et al., 2018), associability, i.e., uncertainty-driven attention (Pietrock et al., 2019), or both (Koenig et al., 2018). Moreover, higher average effects in aversive than appetitive conditioning studies were identified by meta-analysis (Finke et al. 2021), yet it is unclear whether this apparent US-valence effect reflects unsystematic between-study heterogeneity, systematic differences in arousal and/or stimulus intensity, or genuine dissociations in pupil-linked neurocognitive mechanisms. Disentangling these possibilities, and thus establishing whether the pupil provides a unitary or more complex readout, requires well-powered evidence linking varying temporal phases of pupil responses to affective and cognitive processes during learning.

Mechanistically, pupil dilation results from the interplay of two opposing autonomic pathways acting on different timescales: sympathetically mediated activation of the dilator muscle as well as inhibition of parasympathetic outflow to the sphincter (Loewenfeld & Lowenstein, 1993). In line with this, a considerably large body of research has provided evidence for a distinction between rapid parasympathetic and later sympathetic effects unfolding within the same (biphasic) response (e.g., Steinhauer & Hakerem, 1992; Finke et al., 2017; Widmann et al., 2018; Finke et al., 2023a). Similarly, functionally distinct types of pupil responses subserved by partly dissociable brain systems (in addition to the locus coeruleus-noradrenergic [LC-NE] system) and characterized by different latencies and temporal dynamics have been proposed (Strauch et al., 2022). Together, these observations argue in favor of a more fine-grained, temporally resolved approach, rather than averaging over extended windows, extracting peak values, or imposing a canonical response function of fixed shape, which, while useful for characterizing learned fear (PsPM; Korn et al., 2017), may be less suited to capturing multiple, temporally overlapping processes underlying the pupil response (Hoeks & Levelt, 1993; Blini et al., 2024).

Beyond such model-based approaches, state-of-the-art, data-driven techniques to analyze pupil traces include generalized additive mixed modeling (GAMM; Van Rij et al., 2019) and dimensionality reduction via temporal principal component analysis (PCA), which is well-established in EEG research (Dien, 2010) as well as in pupillometry (Siegle et al., 2001; Wetzel et al., 2015; Finke et al., 2017). Both methods are particularly suited to examine complex dynamics in the time course of evoked responses. However, to our knowledge, decomposition of the within-trial temporal structure of pupil dilation, via PCA, has so far not been applied to pupillometry in Pavlovian conditioning, despite recent recommendations regarding its validity (Castellotti et al., 2025) and growing use of single-trial PCA in memory research (e.g., Clewett et al., 2020; Xue et al., 2026, reviewed preprint).

The present study aimed to address both gaps. Specifically, we set out 1. to uncover the temporal composition of pupil dilation during Pavlovian learning; 2. to probe how this temporal signature relates to central constructs of associative learning: the CS+ vs. CS-contrast, the role of US valence, and the interplay of associative strength, associability and prediction error across cue (CS) and outcome (US) onset; and 3. to validate this analysis by examining its convergence with a complementary, assumption-free model of the raw pupil waveform (based on GAMM). To this end, we conducted an individual-participant-data reanalysis (mega-analysis) of primary studies from our lab. Whereas our previous meta-analysis (Finke et al., 2021) pooled aggregate effect sizes - revealing robust differentiation in both early (post-CS) and late (pre-US) response windows, yet marked heterogeneity in window definitions and other methodological choices, alongside a predominance of aversive conditioning studies – it could not establish whether these intervals (and associated differences in effect sizes) reflect distinct underlying processes. By pooling participant-level data, the present mega-analysis instead allows us to dissociate the temporal dynamics of conditioned pupil dilation at the single-trial level. Crucially, all included studies were planned and conducted after that meta-analysis, deliberately harmonized in design and timing to address its limitations as well as further gaps in the literature. In addition, we performed a brief, independent out-of-sample validation as a further generalization test of our main findings. By linking distinct temporal components to specific learning-related variables, this approach promises a more precise functional characterization of conditioned pupil dilation and a firmer basis for its future application as a marker of the processes that drive associative learning.

## 2. Method

### 2.1 Selection of studies, participants

Table 1 lists the major characteristics of all primary studies included in this reanalysis (7 independent samples). All experiments were conducted between 2020 and 2026 at the University of Siegen (Sample ID 1-6) or University of Trier (Sample ID 7), Germany. Datasets from ongoing studies were frozen on June 10, 2026. In contrast to several primary studies included, the present reanalysis was not preregistered. All studies were conducted with approval of the responsible ethics committee (ID 1-6: Ethikrat der Universität Siegen; ID 7: Landesärztekammer Rheinland-Pfalz) and in accordance with the Declaration of Helsinki and involved healthy human individuals only, with no reported intake of psychotropic medication, no mental health problems during the last two years and no evidence of acute or severe, chronic somatic disorders. Cumulative sample size across all experiments amounts to *N* = 385 participants (66.7% women on average, *SD* = 13.5) or *M* = 562 datasets of separate conditioning sessions, respectively. Mean age across all studies, weighted by number of participants, was 23.3 years (*SD* = 4.8). See Table 1 for details on sample characteristics. Exclusion decisions were applied at the level of sessions and trials wherever possible rather than at the participant level. Specifically, we retained all session-level datasets in which participants contributed at least 50% of valid trials per CS category (e.g., CS+_avs_) after quality checks during preprocessing. This strategy minimizes the risk of introducing selection bias into PCA estimation, enhancing the generalizability of the derived component structure. Accordingly, effective sample sizes for the present analysis may deviate slightly from the original *N*s reported in the respective publications.

**Table 1.**
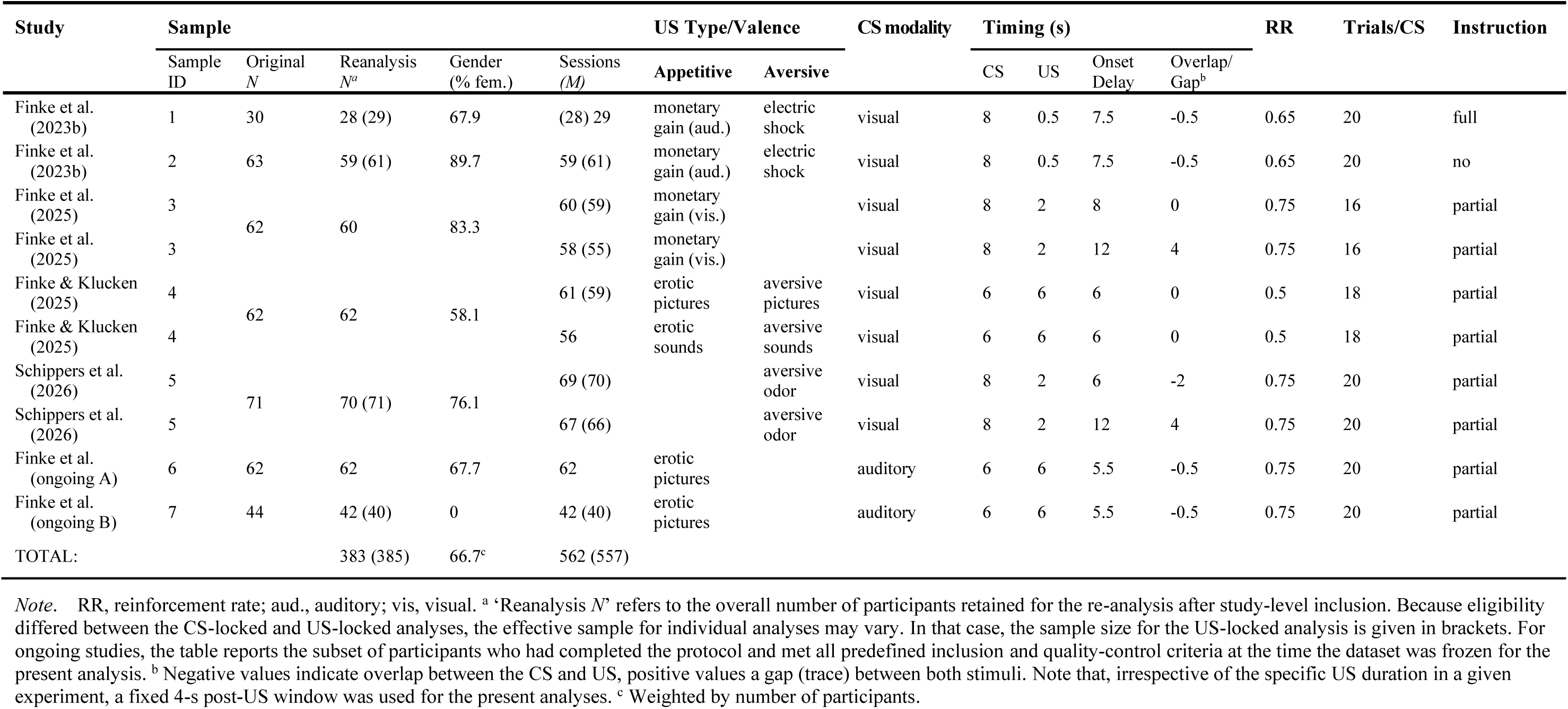
Sample characteristics and design features of the primary studies included.

### 2.2 Design and procedure

#### 2.2.1 General design and procedure

As evident from Table 1, the set of primary studies covered a broad range of variation in US type, valence (aversive vs. appetitive) and duration (range: 0.5-6 s). In contrast, other design features were largely comparable across studies, especially regarding timing characteristics, with CS duration ranging from 6 to 8 s, number of trials per CS from 16 to 20 and reinforcement rate (RR) from 50 to 75%. CS-US onset delay ranged between 5.5 and 12 s, with most studies using a standard delay conditioning paradigm with shortly overlapping or immediately adjacent stimuli and two studies (appetitive: *k* = 1; aversive: *k* = 1) comparing trace and delay conditioning in two separate sessions. Visual CS were used in the majority of studies, with some exceptions (auditory modality: *k* = 2). The general procedure (as described in greater detail in the original publications) was as follows: Participants visited the laboratory on one or (in case of studies involving two conditioning sessions, *k* = 3) on two separate days. Upon arrival at the laboratory on their first visit, they were first informed about the general methods and procedures employed in the study. After signing the informed consent form, they filled in questionnaires about demographic and health-related information (as well as additional questionnaires depending on the study). Then, they were seated at a desk in the testing chamber (in a different room). After the experimenter had attached electrodes for psychophysiological measurements, the eye-tracker was calibrated. Then, participants performed the (first) conditioning session. If there was a second session, it was identically structured (apart from initial instructions, signing of informed consent and questionnaires). Assignment to the order of different sessions (e.g., trace vs. delay) was always counterbalanced between participants. Immediately before each conditioning phase, participants received written instructions on the screen. In most studies, participants were told that they would be presented with a series of (visual or auditory) stimuli which were either followed or not followed by another stimulus (i.e., the respective US), and that they would have to figure out the relationship for themselves (i.e., partial contingency instruction). To assess potential differences between aware vs. unaware learning, two (otherwise identical) experiments (Sample ID 1 and 2) directly compared the effects of a full contingency instruction vs. no instruction. At the end of the last session, participants were thanked and fully debriefed.

#### 2.2.2 Conditioning (acquisition)

During the acquisition phase, each visual CS (i.e., CS+ and CS-) was shown in the middle of the screen. Auditory CS (sine and saw tones varying in frequency) were presented via closed headphones. Assignment to stimuli (stimulus sets) was always counterbalanced. The order of trials was fully randomized, apart from the following constraints: (a) The first trials contained stimuli from all CS conditions; (b) the first CS+ trials were always reinforced; (c) a particular CS did not appear on more than two successive trials. The inter-trial interval (ITI) randomly varied between 11 and 13 s in all studies. In reinforced trials, an US followed the CS at a fixed latency (depending on the specific study, see Table 1). In nonreinforced trials (including all CS-trials), there was either no subsequent stimulus or a neutral, “non-US” control stimulus (e.g., “+ 0.00 €” or neutral picture). During the ITI, including the time before CS onset, a perceptually matched control slide was shown (see Stimuli and apparatus), in order to control for physical influences on pupil size and prevent activation of the light reflex of the pupil.

After acquisition, participants were prompted to provide ratings of subjective arousal, valence and US expectancy for the (respective) CS+ and the CS-(see below).

### 2.3 Stimuli and apparatus

#### 2.3.1 Conditioned stimuli

Greyscale, circular grating stimuli varying either in orientation/rotation angle (Sample ID 1, 2) or in orientation and grid width (within separate sets per session; Sample ID 3, 4) were used as CS in most studies. One study used greyscale pictures of neutral faces (Sample ID 5), and two studies (Sample ID 6, 7) simple tones varying in frequency and wave form. All visual stimuli used as CS were shown in the center of the screen in front of a light-grey background, subtending a visual angle of approximately 5° × 5°. Assignment of stimuli to conditions was always (pseudo-)randomized and counterbalanced between participants.

#### 2.3.2 Unconditioned stimuli

Aversive unconditioned stimuli included: (a) unpleasant electro-cutaneous stimulation (for 0.5 s; Sample ID 1, 2), (b) fear- and disgust-related pictures (Sample ID 4) as well as (c) aversive olfactory stimulation (for 2 s); as appetitive US, (d) a sound of coins (duration: 0.5 s) signaling a monetary win of 1 € (Sample ID 1, 2), (e) a visual verbal cue (text; duration: 2 s) signaling a monetary win of 0.5 € (Sample ID 3), and (f) erotic sound clips (duration: 6 s; Sample ID 4) or (g) erotic pictures of couples (duration: 6 s; Sample ID 4, 6, 7) were employed. Grey-scale pictures equalized for contrast/dispersion and subtending a visual angle of approximately 5° (horizontally) × 3.75° (vertically) were used in Samples 6 and 7, whereas one study (Sample ID 4) used color pictures. A control slide matched to visual CSs (or USs: Sample ID 6, 7) in terms of brightness, contrast and dispersion was presented during the ITI, consisting of a shape (circle, ellipsoid or rectangle, respectively) filled with randomly scattered pixels as well as a small white fixation cross. See the original articles for further technical details on stimulus features and presentation.

#### 2.3.3 Apparatus

In the studies conducted at the University of Siegen (Sample ID 1-6), participants were seated in front of a 27-inch TFT screen (with a resolution of 1920 × 1080 pixels and a refresh rate of 60 Hz) at a viewing distance of 70 cm, with their forehead and chin resting on a head rest mounted on a height-adjustable desk. The experiment was programmed with the Psychtoolbox (Version 3.0.10; Kleiner et al., 2007) implemented in MATLAB (MathWorks, Inc., Natick, MA, USA) and ran on a standard Windows PC. In Trier (Sample ID 7), a TFT screen with a resolution of 1600 × 1080 pixels and a refresh rate of 75 Hz was used (viewing distance: 60 cm), and stimuli were presented using E-Prime 2.0 (PST Software, Inc.).

#### 2.3.4 Acquisition of pupillary data

Pupil diameter was recorded from the right eye at a sampling frequency of 1000 Hz using a video-based infrared eye-tracking device (EyeLink 1000 Plus, SR Research Ltd., Ottawa, Canada) in the Siegen lab (Sample ID 1-6). In Trier (Sample ID 7), a SMI eye-tracker (operating at 500 Hz) was used (iView-X HiSpeed 500, SensoMotoric Instruments, GmbH). Eye-tracker calibration was performed by means of an in-built 9-point calibration routine and repeated until an accuracy of < 1.0° was achieved.

### 2.4 Data recording and reduction

#### 2.4.1 Pupil dilation

Pupillometric data from each study were subjected to the same pre-processing pipeline (using MATLAB, MathWorks, Inc.), applying consistent criteria for artifact correction in accordance with recently published recommendations (Steinhauer et al., 2022) as well as procedures established by prior conditioning studies (Finke et al., 2023b; Korn et al., 2017; Leuchs et al., 2019). After segmentation into long trial epochs spanning from ITI onset through at least 4 s after US onset, data were down-sampled to 100 Hz (by means of a moving window filter selecting the median value across 10-ms epochs). Missing data points (resulting from blink artifacts and small head movements) were replaced by linear interpolation. To harmonize data across samples and paradigms, individual pupil-size time series were z-standardized within participants and sessions. To account for variation in timing features between studies (i.e., CS-US onset delay, etc.; see Table 1) and target responses after both CS and US onset, data were then segmented into CS-locked and US-locked epochs for further analyses (CS-locked analysis: -2 to +6 s post CS onset; US-locked analysis: -2 to +4 s post [expected] US onset). Criteria for trial rejection were applied to both analysis windows separately but consistently: Specifically, epochs containing high amounts (> 50%) of interpolated data or values exceeding a threshold of 3 SDs above or below the participant’s mean pupil size were discarded. For studies with visual CS and/or US, epochs were also discarded if eye gaze was not fixated on the center of the screen (e.g., due to prolonged blinking, distraction etc.) for at least 50% of epoch duration, based on a cut-off window around the participant’s median gaze position (across all trials) covering approximately ± 5° visual angle. Trials with an unreliable pre-stimulus baseline (> 50% missing/interpolated data and/or > 3 *MAD* above or below the intraindividual median of the baseline) were also flagged as invalid. Across datasets (samples/sessions), 8.32% (*SD* = 2.05) of relevant CS epochs and 11.46% (*SD* = 3.70) of US epochs were rejected (excluding completely discarded individual datasets). After data cleaning, pupil traces of each epoch were referenced to a common pre-stimulus baseline by subtracting the mean value (averaged over a period of 500 ms) immediately before CS onset from each subsequent data point. Datasets (sessions) from participants with less than 50% valid trials per CS category and analysis window were excluded from the respective statistical analysis (CS-locked: *m* = 9; US-locked: *m* = 14).

#### 2.4.2 Self-report ratings

Immediately after the acquisition phase, participants were prompted to indicate US expectancy for each CS using a visual analogue scale (VAS) ranging from 0% to 100% (probability of occurrence). Subjective levels of arousal and valence attributed to each CS were assessed via VAS as well, with the anchors ‘calm and relaxed’ to ‘very arousing/tense’ and ‘very unpleasant’ to ‘very pleasant’, respectively. Distances in pixels were converted to arbitrary units (ranging from 0 to 1). Due to technical errors, rating data of some participants were lost.

### 2.5 Signal decomposition and statistical analysis

#### 2.5.1 Extraction of PCA components

After segmentation into trial epochs, baseline-corrected pupil data from both analysis windows (across all studies) were downsampled to 10 Hz, arranged in a wide format and subjected to PCA at the single trial level. Extraction of latent components was performed in *R*, based on the covariance matrix. Due to the high extent of autocorrelation inherent in physiological signals, traditional decisional criteria (such as scree test and parallel analysis) may be considered insufficient. Instead, for each window of analysis, we aimed for a solution that was (a) reproducible across samples (Tucker congruence φs > .95; see below) and (b) jointly accounted for > 80% of total (orthogonal) variance, (c) yielded a ‘simple structure’ (i.e., minimal cross-loadings after Promax rotation, κ = 3; Dien et al., 2010; Wetzel et al., 2015), and (d) remained interpretable, with little temporal overlap between component loadings.

PCA was conducted on the clean dataset, with invalid trials (i.e., trials with high amounts of artifacts or missing data) removed from the signal. To ensure both reliability and generalizability of the final PCA solution, while at the same time avoiding bias due to differences in sample sizes between primary studies as well as selective drop-out resulting from overly strict exclusion criteria, we conducted the PCA based on single-trial data inversely weighted by the number of (valid) trials per participant and the respective sample size of the study (with normalization of weights). This approach ensures that each study and each participant within each study contributes equally to the covariance matrix. Moreover, to get an unbiased PCA structure that captures the within-trial temporal dynamics of associative learning and its effect on pupil size, rather than external sources of arousal, trials containing acoustic startle probes (noise bursts) during the CS interval (Sample ID 3 only) were initially excluded. Loadings of the resulting reference PCA were then used to conduct reliability analyses and to project component scores (indexing the contribution of each component to the pupil signal on each trial) onto all valid trials (including startle trials) for further analyses. (A sensitivity analysis concerning all major outcomes showed that in- or exclusion of startle trials yielded equivalent results.)

#### 2.5.2 PCA stability

The stability/invariance of the component structure (1. within studies, i.e., across experimental trials, as well as 2. between independent studies) was evaluated using Tucker congruence φ (i.e., the uncentered correlation between loadings; Lorenzo-Seva & Ten Berge, 2006). To this end, (1) separate PCAs (with identical number of components) were estimated for each experimental half (median split across trials) across all studies and compared against each other (split-half Tucker congruence). Moreover, (2) PCA solutions with one sample left out (k-1) were iteratively computed and compared with the loading structure based on the respective sample. This ‘leave-one-sample-out’ approach avoids circularity, yielding unbiased estimates in turn (e.g., in contrast to fitting a PCA on the basis of all studies and comparing this reference with each individual study). The range and median of resulting Tucker scores (φ) across all iterations were determined for all candidate solutions (i.e., varying numbers of components). If there were minor differences in the sequence of extraction, highly correlated (i.e., effectively identical) components were matched based on an algorithm as well as visual inspection of loading patterns across time. Scores > .95 signal excellent consistency (Lorenzo-Seva & Ten Berge, 2006), which served as the main criterion for selection. Note that, while solutions based on a higher number of components (as compared to the chosen solution) would all have led to reduced reproducibility in the CS-locked and US-locked analyses (φ_min_ ≤ .8; see Supplementary Material, Table S1), a two-component solution would have been equally stable for the US-locked window. However, as the distinctness and interpretability of the corresponding loading pattern were comparatively lower and congruence would have been strongly diminished with both three (φ_min_ ≤ .85) and five (or more) components (φ_min_ ≤ .55) in this window, we decided in favor of the four-component solution. See Supplementary Material for further information on PCA selection.

#### 2.5.3 Assessment of conditioning effects

For scores of each PCA component, linear mixed-effects models (LMMs) were computed to evaluate which temporal component indexes differential conditioning (i.e., differentiation of responses to the CS as well as unconditioned responding) and whether this differs between appetitive and aversive learning. Models were fitted (with restricted maximum likelihood estimation) using the *lme4* package in R (Bates et al., 2015). In all models, clustering of data was controlled for by entering orthogonal random effects of *Subject* and *Sample* as well as *Sessions* nested in *Subjects*. (Note that models with random slopes for fixed effects [such as *CS Type*] varying by participant were also evaluated but yielded singular fits.) In addition, a core set of major, design-based predictors were always included as fixed effects. For CS-locked analyses, fixed effects comprised *CS Type* with orthogonal, nested contrasts for (1) differential conditioning (CS+ vs. CS-) and (2) *US Valence* (appetitive vs. aversive). In addition to these predictors, responses in the US-locked window were modeled by *US Presence*, again with orthogonal, nested contrasts for (3) unconditioned responding (US+ vs. US-, i.e., absent vs. present, nested within CS+ trials), and (4) *US Valence* (appetitive vs. aversive, nested within US+ trials). Furthermore, to assess influences of other important design-related factors, we performed additional exploratory/sensitivity analyses by separately entering (i) *CS-US Contiguity* (trace vs. delay conditioning) and (ii) *CS Trial* (mean-centered) as well as all respective two-way interactions with *CS Type* and (if applicable) *US Presence*. We refrained from directly comparing effects of the modality of the CS (visual vs. auditory) in the present study, as this design feature was confounded with the type of outcome (i.e., erotic pictures in all primary studies using auditory cues).

To evaluate the significance of fixed effects, initial models were incrementally reduced using likelihood-ratio tests (after refitting based on maximum likelihood estimation). The random-effects structure was kept unchanged across model comparisons. Significance of regression estimates (model coefficients) was assessed by t-tests with Satterthwaite approximation of degrees of freedom. For all analyses, the threshold of significance was set to α = .05, with all *p* values reported as two-tailed and statistical differences at *p* < .10 considered as trends. Since PCA component scores were normalized (with *SD* ≈ 1), all coefficients (*b*) of contrast-coded factors can be interpreted as partially standardized mean differences (analogous to Cohen’s *d*), and we therefore report these coefficients as effect-size indices. Note, however, that underlying *SD*s reflect total-trial variance which is dominated by residual (within-subject) variance and thus directly comparable only to effects in single-trial models rather than Cohen’s *d* based on means aggregated at the subject level (Brysbaert & Stevens, 2018). For the main a-priori contrasts (related to conditioning), we therefore also report Cohen’s *d_z_* (based on aggregated data) to enhance comparability with previous research.

#### 2.5.4 Association with computational learning signals

As a theory-guided characterization, we examined whether the extracted temporal components showed the expected correspondence with learning quantities derived from a hybrid RW-PH model:

(1) *V_t_* = *V_t_*_−1_ + α*_t_*_−1_δ*_t_*_−1_, with

(1a) α*_t_* = γ | δ*_t_*_−1_ | + (1 − γ) α*_t_*_−1_, and

(1b) δ*_t_* = *US_t_* − *V_t_*

To avoid inferential circularity (when examining differential associations with component scores) as well as noisy fits resulting from between-subject and between-study heterogeneity, we refrained from estimating the underlying parameters (constant γ and fixed initial values V_0_, α_0_) based on observed pupil data. Rather, learning quantities were recursively (trial-by-trial) computed based on start values informed by our own as well as independent pupillometric research (Finke et al., 2023b; Pietrock et al., 2019) and used as regressors at the trial level. Accordingly, the model was not intended to provide an optimal description of individual learning behavior, but to generate theoretically motivated, normative regressors against which the functional specificity of the extracted components could be evaluated. Specifically, initial *V* (value/associative strength) was set to 0.5, initial α (associability) to 0.7 and the decay parameter γ (providing higher weight to earlier trials) to 0.1 for each CS (within subjects and sessions). For each component score, a nested series of LMMs was fitted and compared in terms of fit: (1) a null model (random effects only), (2) a model with learning variables only, (3) a combined model with learning variables in addition to design-based contrasts (*CS Type* as well as *US Presence,* if applicable), and (4) a model including the *Learning* × *CS Type* interaction. For analyses of components peaking before or at (expected) US onset, α and *V* were entered as (mean-centered) predictors, indexing antecedent learning states updated on the previous trial. Analyses of components related to outcome responses (after expected US onset) included terms reflecting prediction errors from the current trial, i.e., subsequent learning states. More precisely, to independently model signed (δ) and unsigned prediction errors (|δ|), positive (δ+) and negative (δ-) values were used as separate regressors, indexing unexpected reinforcement vs. omission. If there was evidence for an overall significant association with learning signals (based on lower AIC and higher likelihood ratio relative to the null model), the incremental significance of each individual predictor (e.g., α) was evaluated independently. To this end, effects with significant coefficients were directly assessed by comparing a design model with and without this learning variable. To ensure interpretability of model coefficients, multicollinearity was checked based on *VIF* scores for all fitted models (to avoid models with strongly collinear predictors).

Furthermore, to assess robustness against the choice of fixed starting values, we performed additional sensitivity analyses (reported in the Supplementary Material) by iteratively computing LMMs with learning signals as predictors and each component as dependent variable for a grid of plausible fixed values. For each run, standardized effect estimates (β) were calculated and their distribution summarized across the grid. Across all combinations of parameters, the pattern of coefficients remained highly consistent with the reference solution, indicating that the reported associations between component scores and learning quantities do not depend on the specific set of parameters chosen. See the Supplementary Material for further details on this analysis.

#### 2.5.5 Association with self-report learning indices

Furthermore, to link variation in self-reported (‘subjective’) aspects of learning to different pupillary response components, scores of each pre-US component were regressed on self-report ratings of *US Expectancy* (of the respective outcome), *Valence* and *Arousal*, based on data from all participants for whom post-acquisition CS ratings were available (*n* = 358). Again, we compared models containing only these variables as fixed effects against null models and models with additional, design-based predictors (see above). For descriptive purposes, we also calculated single-predictor models. Note that we refrained from analyzing the relationship of post-US component scores and self-report ratings because post-acquisition ratings indicated CS-related arousal and valence, which makes any straightforward interpretation with US-evoked responses difficult (ratings of US valence and arousal were only available for a subset of studies).

#### 2.5.6 Triangulation based on generalized additive mixed modeling

Using the *mgcv* and *itsadug* packages in *R* (Wood et al., 2018; Van Rij et al., 2015), generalized additive mixed modeling (GAMM) of average pupillary waveforms (Van Rij et al., 2019) and their interplay with experimental factors was applied, based on 10-Hz trial-level data. GAMMs estimate nonlinear relationships through penalized regression splines (smooth terms), allowing to fit the time course of evoked responses both within and between trials without a-priori assumption about its specific shape. For the present data, the average shape of waveforms within trials (*Time*), separately for CS-locked and US-locked responses, was modeled as marginal smooths. To assess whether this shape was modulated by *CS Type* (entered as an ordered factor), independently penalized difference smooths (by CS+_app_, CS+_avs_, with CS-as reference) were estimated. For US-locked analyses, effects of *US Presence* (in addition *to CS Type*) were modeled as difference smooths relative to non-reinforced trials (separately for aversive and appetitive outcomes). Moreover, parametric terms for contrasts of *CS Type* (and *US Presence,* respectively) as well as random intercepts analogous to the LMM analysis were entered. In addition, to improve overall model fit, a smooth for change across trials (e.g., habituation) and a random smooth for *Time* (per participant) were fitted (thereby accounting for unsystematic between-subjects variability in waveforms). Also, we adjusted for *CS-US contiguity* (trace vs. delay) and *Startle Probe* (present vs. absent) as ‘nuisance’ difference smooths (including corresponding parametric terms). To control for autocorrelation of adjacent samples within trials, a first-order autoregressive error parameter was estimated and accounted for (ρ = .97), with the onset of each trial marked. The *bam* function with fast restricted maximum likelihood estimation was used for model fitting. To avoid overfitting, the complexity of smooths (‘wiggliness’) was adapted based on both statistical diagnostic criteria (reflecting the proportion of residual variance of adjacent samples, i.e., k-index; Wood et al., 2016) and visual inspection (comparison of model-predicted values and aggregated raw data). To localize effects across *Time*, simultaneous confidence bands were computed for each major contrast. The posterior distribution of each difference was approximated by drawing 5000 samples from the multivariate normal and calculating the maximum absolute studentized deviation across all data points. The 95^th^ percentile of the resulting distribution of maxima served as critical value for confidence bands, controlling for family-wise error rate across the whole time series (adjacent regions with the band excluding zero were treated as a significant cluster). Furthermore, the correspondence between PCA-reconstructed waveforms (grand averages of difference waves for the CS+ vs. CS-contrast and the effect of *US Presence*, for each level of *US Valence*) and GAMM-predicted waveforms was evaluated by means of Pearson correlations.

#### 2.5.7 Robustness checks and out-of-sample validation

To assess the robustness of our principal findings to specific analytical choices, we conducted a series of targeted sensitivity analyses concerning PCA specification, stability across study samples, model parameterization, and the analytical framework used to characterize the pupillary waveform. These analyses are described in the respective sections above as well as the Supplementary Material. Moreover, to assess generalizability beyond the originating research group (as suggested during peer review), we additionally analyzed publicly available raw pupillometry data from seven independent conditioning experiments (Korn et al., 2017; Xia et al., 2025). After data preprocessing (as outlined above), we first independently estimated the PCA solutions (as specified in our primary analysis) in the external data and quantified their correspondence with the primary solutions using Tucker congruence. For functional validation, the primary PCA scoring solution was kept fixed and projected onto all valid trials in the external dataset, after which the same core design-based contrasts were tested using LMMs. A more detailed description is provided in the Supplementary Material.

## 3. Results

### 3.1 Signal decomposition

#### 3.1.1 CS-locked analysis

Based on all studies, four components were extracted in the CS-locked window, explaining 88% of total variance (before rotation). The time course of loadings after promax rotation is illustrated in Figure 1 (upper panel, left), showing an early, ‘Pre-CS’ component (with a peak around 2 s prior to CS onset), another component peaking shortly after CS onset (‘Post-CS1’: +1.6 s), and two later components, each with similar variance after rotation (‘Post-CS2’: +3.3 s; ‘Post-CS3’: +5.5 s). The chosen solution was highly stable across acquisition halves (Tucker congruence: φs > .99) and did not vary between primary studies (median φ_min_ = 1; range: [.990 1]), showing excellent consistency. See Method and Supplementary Material for further PCA details.

**Figure 1.**
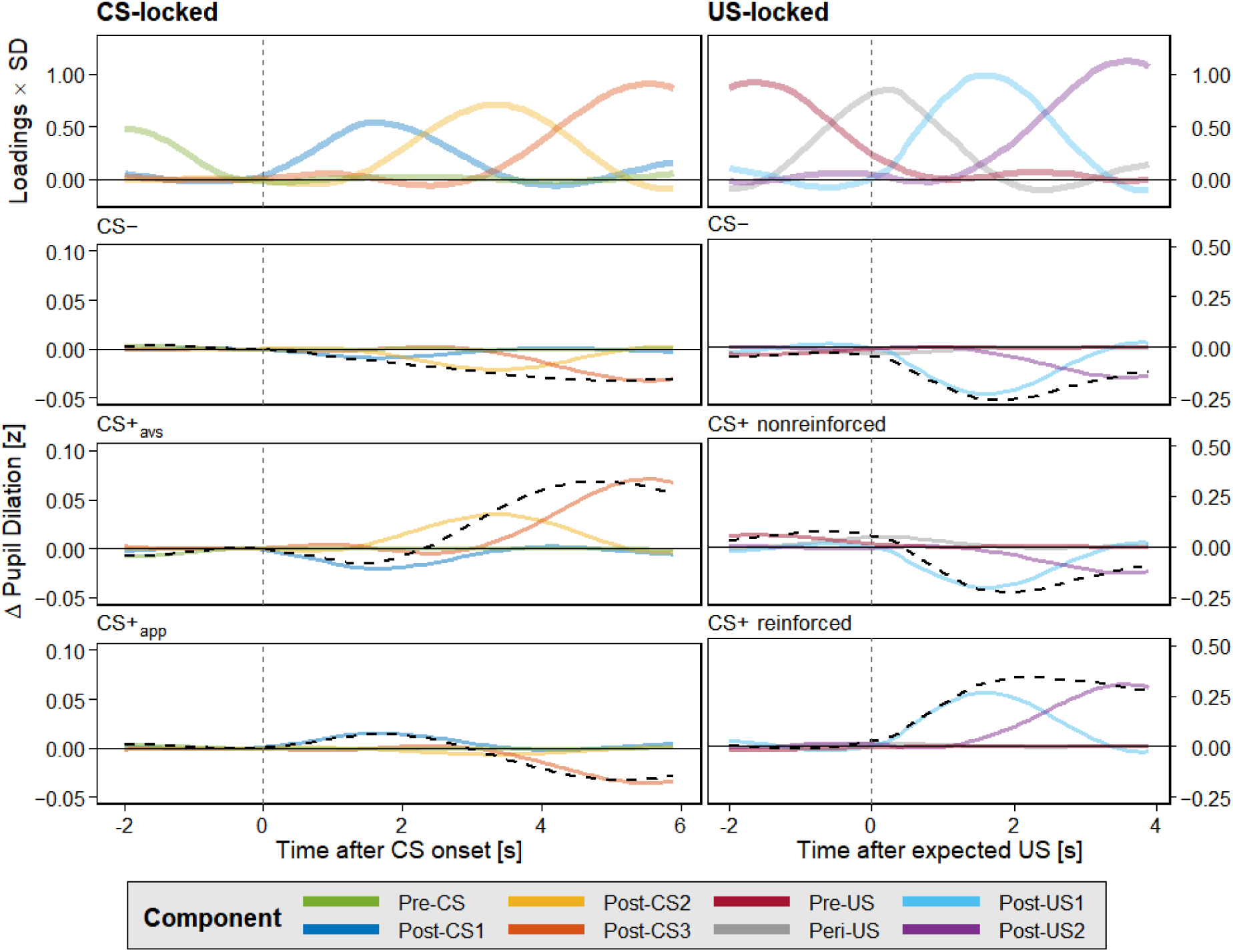
Temporal PCA results. Upper panel: Component loadings (scaled by their respective SD). Panels below: condition-specific contributions (PCA reconstruction from condition-specific component scores), shown separately for the CS- and US-locked analysis. *Note*. CS+_avs_: aversive conditioned stimulus; CS+_app_: appetitive conditioned stimulus; CS-: never reinforced control stimulus.

As shown in Figure 1 (left panels), the extent of differentiation between CS+ and CS-varied (descriptively) between aversive and appetitive cues as well as early and late components. Specifically, the early (Post-CS1) component showed a relatively higher affinity to appetitive conditioning, and vice versa (see 3.2, below).

#### 3.1.2 US-locked analysis

In the US-locked window, four components emerged that together explained 94% of total variance. The temporal pattern fits with two main components reflecting unconditioned responding: one shortly after US onset, with similar latency as in the CS-locked window (‘Post-US1’: +1.6 s) and a later one towards the end of the window (‘Post-US2’: +3.6 s). By contrast, a third component peaked markedly before onset (‘Pre-US’: -1.6 s), while the loadings of the fourth, slightly smaller component showed a steady increase until very shortly after US onset (‘Peri-US’: +0.2 s). See Figure 1 (upper panel, right) for the pattern of loadings after promax rotation. Again, this component structure was largely rotation-invariant, highly stable across acquisition halves (Tucker congruence: φs ≥ .98), and did not vary between primary studies (median φ_min_ = .980; range: [.960 .990]). See Method and Supplementary Material for further PCA details.

The pattern of condition-specific contributions to each component (see Figure 1, right panels) supports the view that the two (temporally) earlier components largely capture anticipatory (conditioned) responses, while both components peaking after US onset reflect unconditioned (and possibly also omission-related) responses.

### 3.2 Conditioning effects

#### 3.2.1 CS-locked analysis

As reported in Table 2, scores of all post-CS components were significantly modulated by *CS Type*, indicating differential conditioning of pupil size in a broad window after cue onset. All respective models explained significantly more variance than a random-effects-only model (all χ^2^s[2] > 9.3, *p*s ≤ .010). Notably, the global contrast between the CS+ and CS-(Post-CS2: *d_z_* = 0.19 [0.09, 0.29]; Post-CS3: *d_z_* = 0.15 [0.05, 0.25]) was not significant for the temporally first (Post-CS1) component, whereas the *US Valence* contrast was. Follow-up tests (with Bonferroni-Holm correction) revealed that differential responses (relative to the CS-) in this initial phase were preferentially expressed in appetitive learning (*p* = .003; *d_z_* = 0.19 [0.08, 0.30]), whereas early responding to aversive cues was characterized by a (non-significantly) reversed pattern (*p* = .119). The difference between both valence categories (aversive vs. appetitive CS+) was significant as well (*p* < .001). There was no reliable statistical evidence for an interplay of *CS Type* with *US Valence* for the two later components (within the CS-locked window), presumably marked by more general response differentiation. As expected, no effects on the Pre-CS component emerged (all *p*s > .21), i.e., during pre-stimulus baseline, which was not favored over a null model (χ^2^[2] = 2.54; *p* = .28).

**Table 2.**
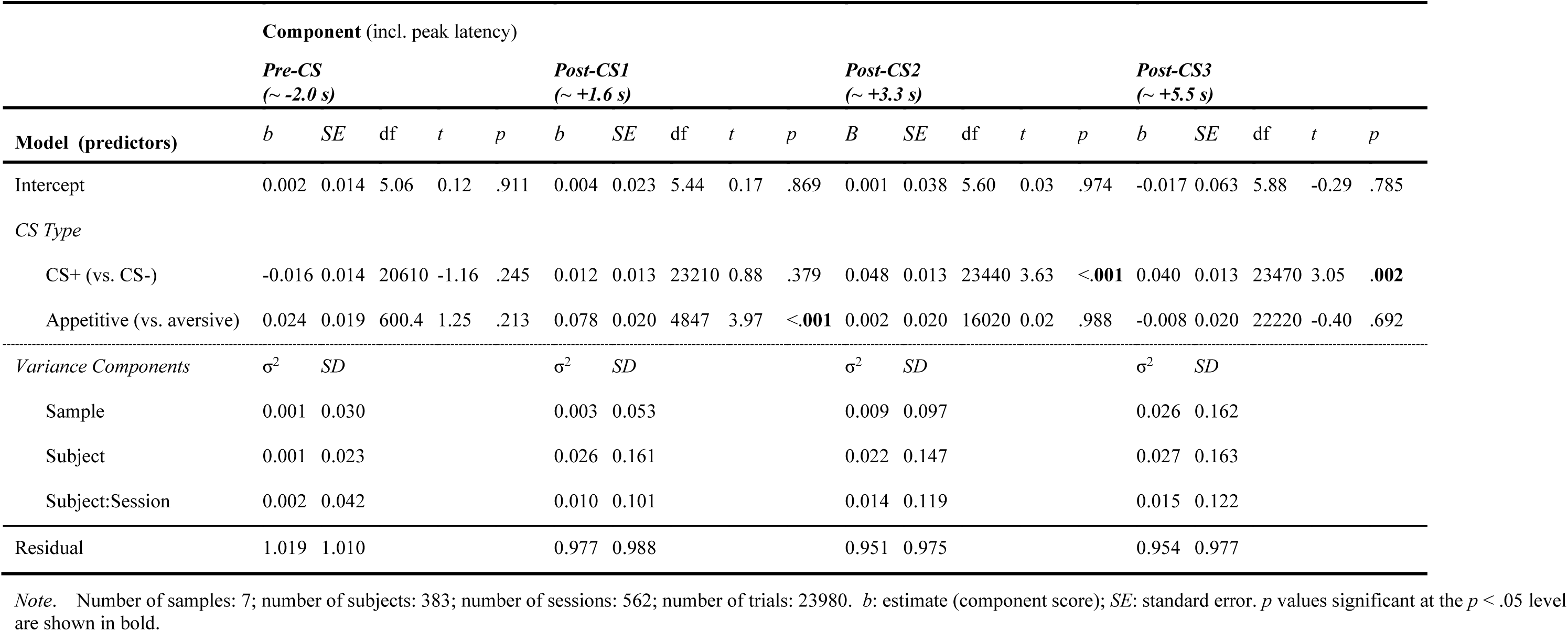
Model parameters of primary linear mixed-effects models predicting component scores in the CS-locked analysis window.

##### Further design-based predictors

Importantly, no evidence for any effects or interactions involving *CS-US Contiguity* emerged for any CS-locked component (all *p*s ≥ .18). The effect of *CS Trial* was significant for the early and middle components (Post-CS1: *p* = .013; Post-CS2: *p* = .019), indicating decreasing response levels over time (i.e., habituation), irrespective of *CS Type* (*p*s ≥ .31). See Supplementary Material for detailed results.

#### 3.2.2 US-locked analysis

In the analysis of US-locked component scores, a clear dissociation between effects on Pre-US scores vs. subsequent components emerged, with tentative evidence for general (valence-unspecific) CS+ vs. CS-discrimination before US onset and a combination of valence-specific and outcome-driven modulation in the Post-US window (see Table 3). All models except Pre-US (modeled with the same fixed-effects structure as outcome-related components; χ^2^[4] = 8.57, *p* = .073) explained significantly more variance than the respective null model (all other χ^2^s[4] > 43.6, *p*s < .001). Since effects of US delivery were statistically absent before US onset (as expected), the only reliable effect on Pre-US scores was a general *CS Type* contrast (*p* = .042; *d_z_* = 0.12 [0.02, 0.22]). In the window after (expected) US onset, pupil size was higher in CS+ trials overall (both Post-US1 and Post-US2; see Table 3), which was initially (Post-US1) more strongly expressed after aversive than appetitive cues. However, outcome-related effects (of *US Presence*) on Post-US1 (*d_z_* = 0.98 [0.86, 1.10]) and Post-US2 (*d_z_* = 0.92 [0.81, 1.04]) scores did not differ by *US Valence* at first (Post-US1), but became slightly more pronounced for appetitive outcomes during later stages of responding.

**Table 3.** Model parameters of primary linear mixed-effects models predicting component scores in the US-locked analysis window.

| Model (predictors) | Component (incl. peak latency) |  |  |  |  |  |  |  |  |  |  |  |  |  |  |  |  |  |  |  |
| --- | --- | --- | --- | --- | --- | --- | --- | --- | --- | --- | --- | --- | --- | --- | --- | --- | --- | --- | --- | --- |
|  | *Pre-US*<br>(~ -1.6 s) |  |  |  |  | *Peri-US*<br>(~ +0.2 s) |  |  |  |  | *Post-US1*<br>(~ +1.6 s) |  |  |  |  | *Post-US2*<br>(~ +3.6 s) |  |  |  |  |
|  | *b* | *SE* | df | *t* | *P* | *b* | *SE* | df | *t* | *p* | *b* | *SE* | df | *t* | *p* | *b* | *SE* | df | *t* | *p* |
| Intercept | -0.007 | 0.051 | 5.88 | -0.13 | .902 | 0.017 | 0.066 | 5.89 | 0.26 | .801 | -0.070 | 0.092 | 6.07 | -0.76 | .474 | -0.024 | 0.049 | 5.83 | -0.49 | .640 |
| *CS Type* |  |  |  |  |  |  |  |  |  |  |  |  |  |  |  |  |  |  |  |  |
| CS+ (vs. CS-) | 0.037 | 0.014 | 22660 | 2.71 | **.007** | 0.059 | 0.014 | 22700 | 4.35 | **<.001** | 0.250 | 0.013 | 22650 | 19.54 | **<.001** | 0.203 | 0.013 | 22660 | 15.22 | **<.001** |
| Appetitive (vs. aversive) | -0.009 | 0.020 | 20670 | -0.42 | .672 | -0.074 | 0.020 | 21960 | -3.65 | **<.001** | -0.074 | 0.019 | 22530 | -3.90 | **<.001** | 0.017 | 0.020 | 20480 | 0.86 | .391 |
| *US Presence* |  |  |  |  |  |  |  |  |  |  |  |  |  |  |  |  |  |  |  |  |
| US+ (vs. US-) | -0.021 | 0.018 | 22660 | -1.18 | .237 | 0.021 | 0.018 | 22690 | 1.16 | .248 | 0.488 | 0.017 | 22650 | 29.05 | **<.001** | 0.396 | 0.017 | 22660 | 22.72 | **<.001** |
| Appetitive (vs. aversive) | -0.024 | 0.036 | 22660 | -0.67 | .501 | -0.097 | 0.035 | 22690 | -2.75 | **.006** | 0.010 | 0.033 | 22640 | 0.30 | .763 | 0.081 | 0.034 | 22660 | 2.34 | **.019** |
| *Variance Components* |  |  |  |  |  |  |  |  |  |  |  |  |  |  |  |  |  |  |  |  |
| | $\sigma^2$ | *SD* | | | | $\sigma^2$ | *SD* | | | | $\sigma^2$ | *SD* | | | | $\sigma^2$ | *SD* | | | |
| Sample | 0.017 | 0.132 |  |  |  | 0.029 | 0.171 |  |  |  | 0.058 | 0.240 |  |  |  | 0.016 | 0.125 |  |  |  |
| Subject | 0.016 | 0.128 |  |  |  | 0.019 | 0.139 |  |  |  | 0.036 | 0.191 |  |  |  | 0.017 | 0.129 |  |  |  |
| Subject:Session | 0.030 | 0.172 |  |  |  | 0.011 | 0.106 |  |  |  | 0.039 | 0.198 |  |  |  | 0.044 | 0.209 |  |  |  |
| Residual | 0.960 | 0.980 |  |  |  | 0.945 | 0.972 |  |  |  | 0.836 | 0.915 |  |  |  | 0.903 | 0.950 |  |  |  |

*Note.* Number of samples: 7; number of subjects: 385; number of sessions: 557; number of trials: 23181. *b*: estimate (component score); *SE*: standard error. *p* values significant at the $p < .05$ level are shown in bold.

Notably, the Peri-US component, bridging CS and US presentation, was sensitive to valence information as well: In contrast to the preceding Pre-US component, responses were significantly larger after presentation of aversive (than appetitive) CS+ (*p* < .001) and differential responding (relative to the CS-) was significant only in aversive (*p* < .001; *d_z_* = 0.40 [0.26, 0.53]) but not in appetitive CS+ trials (*p* = .163 [corrected]; *d_z_* = 0.10 [-0.01, 0.21]). Given this component’s temporal location and the significant *US Valence × US Presence* interaction, we performed further post-hoc analyses to rule out that it was solely driven by delivery of aversive outcomes. Importantly, estimating the CS contrast for non-reinforced CS+ trials only, response differentiation remained significant for aversive (*b* = 0.061, *SE* = 0.024, *z* = 2.52, *p* = .035 [corrected]) but not appetitive conditioning (*b* = 0.036, *SE* = 0.022, *z* = 1.69, *p* = .181) even when no US was delivered. Contributions of differential responding to aversive cues were thus present on all trials (rather than reinforced trials only) and comparable to the increment added by aversive outcomes (*b* = 0.069). Taken together, this indicates that both anticipatory responses and outcome evaluation contributed to Peri-US scores and that these preferentially reflect aversive learning (with substantially weaker differentiation after appetitive CS+).

##### Further design-based predictors

For the Pre-US and Peri-US components, significant effects of *CS-US Contiguity* emerged (*p*s < .001), indicating overall reduced levels of pupil dilation at longer outcome delays. Importantly, differences between responses in trace vs. delay conditioning were again not significantly modulated by *CS Type* (all *p*s > .44). Moreover, no effects of *CS Trial* or *CS Trial* × *CS Type* approached significance for these components (all *p*s > .26). By contrast, the *CS-US Contiguity* × *US Presence* interaction attained significance for components in the post-US window (*p*s < .001), indicating substantially blunted responding to outcomes in trace conditioning (Post-US1: *b* = -0.491, *SE* = 0.035, *t*[22680] = -13.98, *p* < .001; Post-US2: *b* = -0.277, *SE* = 0.036, *t*[22700] = -7.56, *p* < .001). Early (Post-US1), but not late responses in this window decreased over time (*p* < .001), but the slope of this change did not significantly differ between reinforced and non-reinforced trials, suggesting little specific habituation. See Supplementary Material for more information.

Since the (CS-locked) Post-CS3 and (US-locked) Pre-US components showed a very similar profile overall (see also 3.3 and 3.4, below), and in fact overlap in several (delay conditioning) studies with shorter CS-US onset delays, we performed an additional exploratory analysis to test this assumption. To this end, CS-locked and US-locked trial data from all delay conditioning datasets (sessions) were matched based on subject and trial ID. Confirming our interpretation, the correlation between scores of both components (across trials/participants) was *r* = .88, and all effects of experimental conditions were effectively abolished (i.e., near zero) when controlling for the respective other component as covariate.

### 3.3 Computational learning signals

#### 3.3.1 Antecedent learning signals (post-CS)

Across all component scores peaking before (or around) US onset, model-derived antecedent learning variables improved fit over the respective null model (see Table 4, A), indicating systematic, theory-consistent associations with trial-by-trial value (*V*) and associability (α). This association was most evident for Post-CS1 scores, i.e., early responses, as suggested by a *Learning* × *CS Type* interaction. Collinearity was moderate (*VIF* of interaction terms < 4). Inspection of model coefficients revealed that both α and *V* jointly predicted Post-CS1 scores (incrementally explaining variance over a purely design-based model). While the α × *CS Type* interaction was non-significant (overall effect of α across cues: *b* = 0.161, *SE* = 0.058, *t*[23540] = 2.76, *p* = .006), *V* gated responses specifically for reinforced cues (i.e., CS+; *b* = 0.082, *SE* = 0.031, *z* = 2.67, *p* = .008; irrespective of *US Valence; p* = .754), whereas the direction for the CS-was reversed (*b* = -.268). With the appetitive-aversive contrast still significant (*p* = .001), this pattern suggests that both a graded value signal and a dynamic learning rate (scaled by prediction errors from previous trials) may contribute incrementally to responses in this early window (see Figure 2A for illustration).

**Figure 2.**
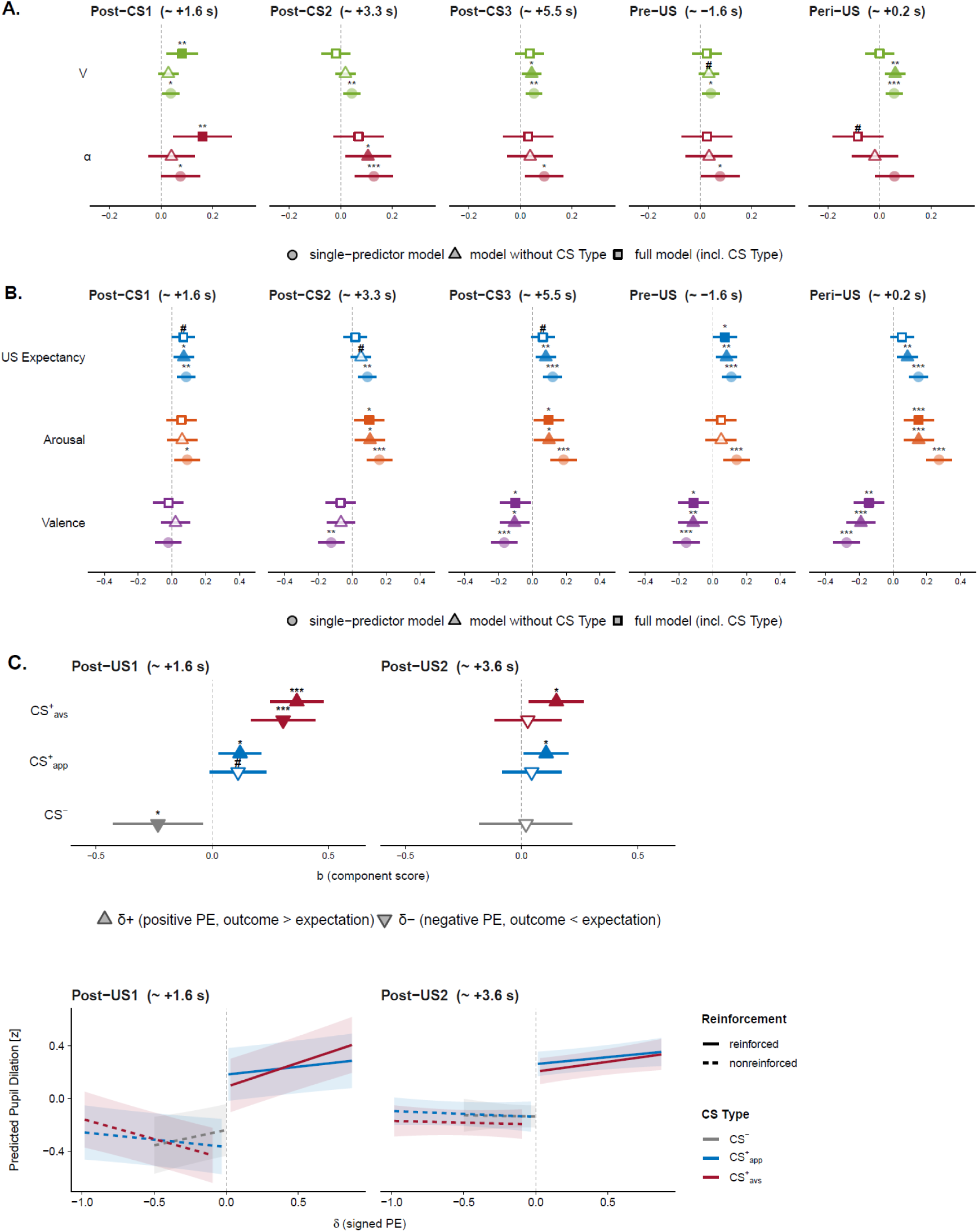
Associations (based on linear mixed-effects models) of A. antecedent learning signals (value, *V*; associability, α) and B. self-report learning indices (*US Expectancy*, *Arousal* and *Valence* ratings) with conditioned pupil dilation across PCA components (before/around US onset). *Note.* X-axis: *b* estimate (component score). Squares: estimates based on full models (A. model 3 [cf. Table 4]: *score* ∼ *CS_Type* + *V* + α + random effects; note that the coefficient shown for Post-CS1/*V* is based on the pooled effect for the CS+, due to the significant *V* × *CS Type* interaction; B. model 6: *score* ∼ *CS_Type* + *US_Expectancy* + *Arousal* + *Valence* + random effects); triangles: estimates based on models without design-based predictors (A. model 2: *score* ∼ *V* + α + random effects; B. model 5: *score* ∼ *US_Expectancy* + *Arousal* + *Valence* + random effects); circles: simple mixed-model regression estimates for each predictor. Error bars indicate 95% CI. C. Differential associations (by levels of *CS Type*) with positive vs. negative prediction errors (PE) for response components after (expected) US onset. *Note.* CS+_avs_: aversive conditioned stimulus; CS+_app_: appetitive conditioned stimulus; CS-: never reinforced control stimulus. The negative prediction error (δ-) was rectified for ease of interpretation. Error bars indicate 95% CI.

**Table 4.**
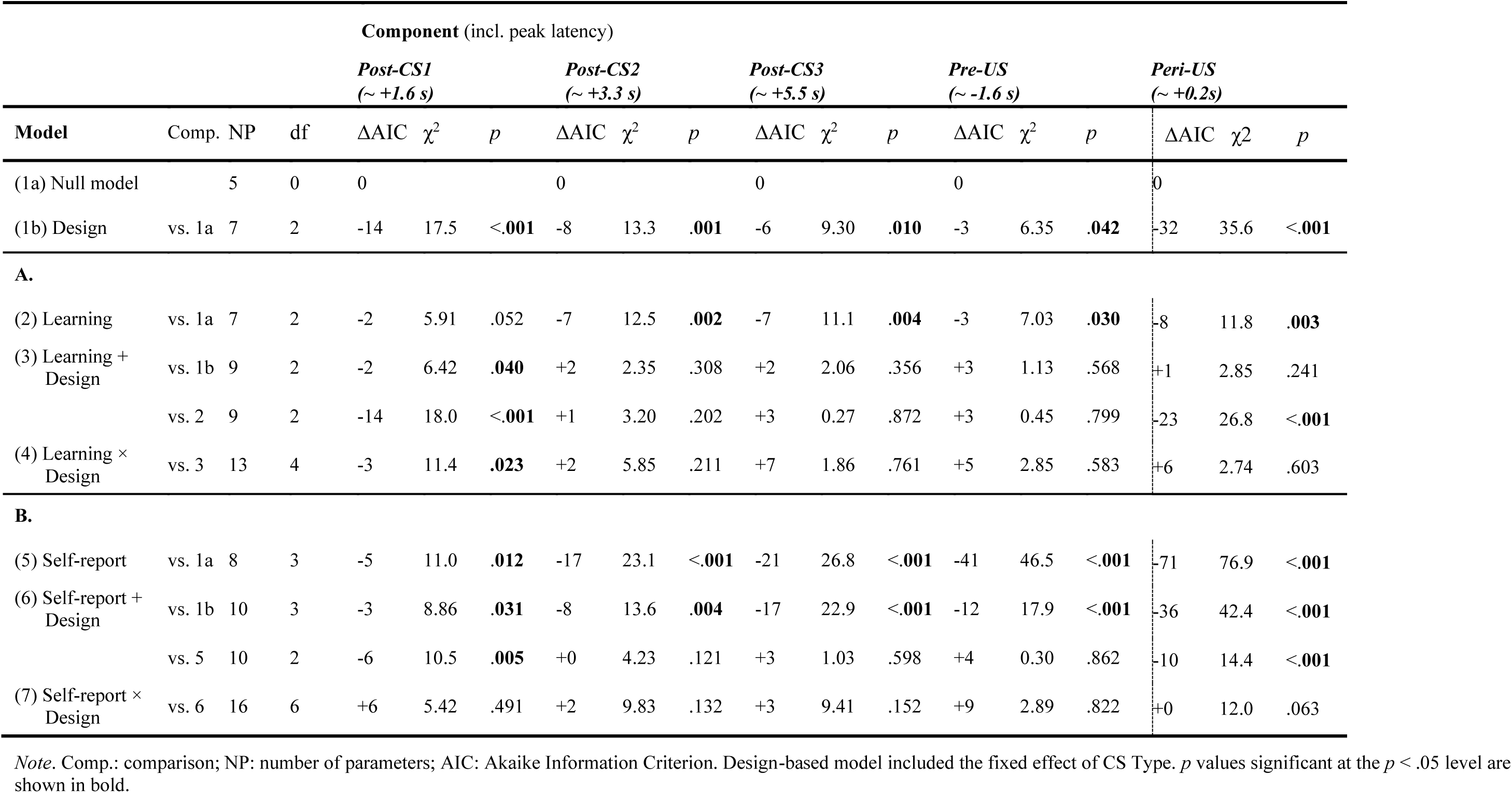
Comparison (goodness of fit) of models based on A. antecedent learning signals (*V*, α) and B. self-report learning indices for post-CS components.

By contrast, learning variables did not reliably contribute beyond design-based predictors for the remaining post-CS components: Inspection of model coefficients (see Figure 2, A: triangle symbols) revealed component-specific links to α (Post-CS2: *p* = .018; negative trend for Peri-US: *p* = .092) vs. *V* (Post-CS3: *p* = .025; marginal for Pre-US: *p* = .088; Peri-US: *p* = .002; all other *p*s > .35), but none remained reliable after adjustment for *CS Type* (i.e., reinforcement/valence of cues; all χ^2^s[1] < 2.9, *p*s > .24; see Figure 2, A: square symbols). Although these later components were thus consistent with hybrid (RW-PH) learning dynamics and even suggested dissociable temporal signatures (differential associations with α vs. *V*), they did not reliably explain variance in pupil responses beyond design-based predictors, as modeled expected value approaches the manipulated contingencies.

#### 3.3.2 Prediction-error-related learning signals (post-US)

As anticipated, post-US components showed clear evidence of specific, incremental contributions of learning signals (i.e., prediction errors) to pupil-size modulation (see Table 5): Inspection of model coefficients indicated that the associations with the earlier component (Post-US1) showed a pattern consistent with unsigned prediction error (|δ|) signaling, i.e., increased responses to both unexpected outcomes and omissions. Collapsing across positive and negative prediction errors, this effect was significantly (*p* < .001 [Bonferroni-Holm corrected]) more pronounced in aversive (*b* = 0.320, *SE* = 0.042, *t*[22730] = 7.62, *p* < .001) than appetitive learning (*b* = 0.106, *SE* = 0.036, *t*[22680] = 2.97, *p* = .003), indicating stronger prediction-error-evoked responses in the early post-US window after expectation of aversive outcomes. In contrast, the later component (Post-US2) tracked positive prediction errors only, i.e., expectation violations in terms of unexpected (appetitive as well as aversive) outcomes (pooled effect: *b* = 0.120, *SE* = 0.037, *t*[22680] = 3.26, *p* = .001), apparently irrespective of valence (see Table 5, model 4). See Figure 2 (C) for illustration. A sensitivity analysis which additionally included the interaction with *US Presence* confirmed that prediction-error-related effects were indeed additive to effects of reinforcement (χ^2^[1] = 0.77, *p* = .38; see Figure 2, C, lower panel).

**Table 5.** Comparison (goodness of fit) of models based on prediction-error-related learning signals (δ+, δ-) for post-US components.

| Model | Comp. | NP | df | Component (incl. peak latency) |  |  |  |  |  |
| --- | --- | --- | --- | --- | --- | --- | --- | --- | --- |
|  |  |  |  | *Post-US1*<br>(~ +1.6 s) |  |  | *Post-US2*<br>(~ +3.6 s) |  |  |
| | | | | ΔAIC | $\chi^2$ | *p* | ΔAIC | $\chi^2$ | *p* |
| (1a) Null model |  | 5 | 0 | 0 |  |  | 0 |  |  |
| (1b) Design | vs. 1a | 9 | 4 | -1501 | 1509 | <.001 | -960 | 967.8 | <.001 |
| (2) Learning | vs. 1a | 7 | 2 | -1020 | 1024 | <.001 | -393 | 610.6 | <.001 |
| (3) Learning +<br>Design | vs. 1b | 11 | 2 | -41 | 44.5 | <.001 | -7 | 11.3 | .004 |
|  | vs. 2 | 11 | 4 | -522 | 529.9 | <.001 | -360 | 368.5 | <.001 |
| (4) Learning ×<br>Design | vs. 3 | 14 | 3 | -21 | 27.5 | <.001 | +5 | 0.84 | .840 |

Note. Comp.: comparison; NP: number of parameters; AIC: Akaike Information Criterion. Design-based model included the fixed effect of CS Type as well as US Presence. *p* values significant at the $p < .05$ level are shown in bold.

Importantly, these associations were robust to the choice of fixed RW-PH starting values, i.e., stable across a broad grid of combinations of *V*_0_, α_0_ and γ (see Supplementary Material, Table S4 and Figure S6). The valence-specific modulation of prediction-error responses in the post-US window was particularly stable.

### 3.4 Associations with self-report learning indices

Overall, self-report indices of conditioning, reflecting both inter- and intraindividual variation in the perception of cues following acquisition, were significantly related to pupil dilation across all CS-related PCA components, as evident from comparisons with null models (see Table 4, B). Moreover, inclusion of self-report indices improved model fit over a design-based model for all components peaking before/around US onset, indicating incrementally explained variance.

As there is a high natural overlap between *CS Type* and *US Expectancy* (which essentially reflects the contrast between reinforced CS+ and non-reinforced CS-), it is unsurprising that controlling for this design factor substantially reduces associations with expectancy ratings. Therefore, and since no interaction with *CS Type* attained significance, our interpretation focuses on models retaining fixed effects of *US Expectancy*, *Arousal* and *Valence* only. Model coefficients (see Figure 2, B: triangle symbols) suggest a consistent, yet differential pattern of associations with temporal response phases: While early responses significantly predicted *US Expectancy* only (when controlling for intercorrelations in a single model), associations with *Arousal* become more robust as the temporal position shifts toward US onset (especially for the Peri-US component). Similarly, for valence (ranging from unpleasant to pleasant) substantial negative correlations with late (yet not early) components emerged, which again became largest and most reliable around US onset. Together, this aligns with the pattern of design-based (valence-specific) contrasts reported above.

### 3.5 Generalized additive mixed modeling of pupil dilation responses

To corroborate the findings from PCA-based LMM analyses, generalized additive mixed models (GAMMs) were additionally computed, providing largely converging evidence for (a) robust differential conditioned responses across both levels of *US Valence*, starting from about 2.4 s after CS onset (in aversive conditioning), as well as (b) a partial valence-dependent dissociation of response phases, with significant early CS+ vs. CS-discrimination (from ca. +0.5 s onward) in appetitive (but not aversive) learning (see Figure 3). Accordingly, the CS+ vs. CS-contrast differed significantly between both valence categories from 0.4 to 1.9 s in the CS-locked window. In line with LMM results, there was also evidence for a short early cluster (ca. 0.5-0.9 s) characterized by significant, inverse response differentiation for the aversive CS contrast (i.e., CS+_avs_ < CS-). Furthermore, (c) GAMM-predicted responses in aversive US+ trials were larger in a broad early cluster (ranging from around 1.2 s before to 1.7 s after US onset). As illustrated in Figure 3, this temporal dynamic corresponds to the transient overlap between the (partly anticipatory) Peri-US and the outcome-locked Post-US1 components. By contrast, later clusters showed the opposite pattern (indexing increased sustained dilation to appetitive outcomes). Consistent with the prediction-error analysis, responsivity in non-reinforced (US-) trials (US omissions) was significantly larger after aversive cues (starting around 0.7 s after US onset), which was aligned with selective modulation of the Post-US1, but not Post-US2, component. Notably, intercorrelations between marginal waveforms predicted by GAMM, waveforms reconstructed from PCA and empirical grand averages in each condition were consistently high (all *r*s ≥ .86; see Supplementary Figure S7). Correlations between GAMM- and PCA-based difference waves (CS+ vs. CS-) were likewise high overall, but more consistent in aversive conditioning. For details on final GAMMs (model coefficients, etc.) see Supplementary Material (Table S5).

**Figure 3.**
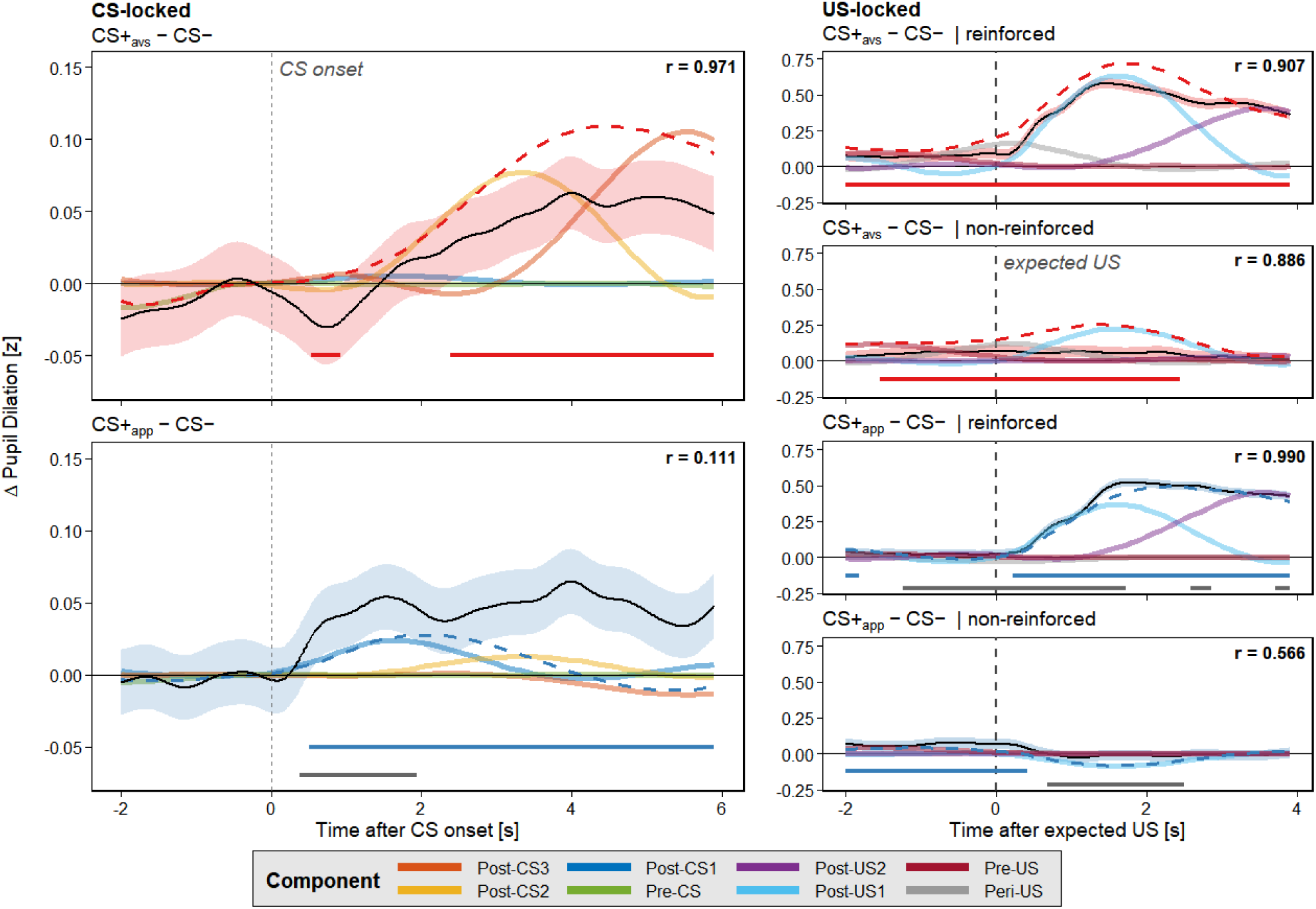
Overlay of (1) PCA-based and (2) GAMM-based reconstructed difference waves (CS+ vs. CS-), split by CS-locked (left) and US-locked (right) windows as well as reinforced vs. non-reinforced trials (upper panels: CS+_avs_ [aversive] vs. CS-; lower panels: CS+_app_ [appetitive] vs. CS-). (1) Colored lines: component contributions; dashed lines: overall waveform based on PCA. (2) Solid lines with shaded bands (95% CI): GAMM-derived conditional means. *r* values indicate Pearson correlations between PCA-based and GAMM-based waveforms in each condition. Colored bars at the bottom indicate significant differences in conditional means, i.e., differential responses (CS+ vs. CS-), based on simultaneous 95% confidence intervals; likewise, grey bars indicate significant differences for the contrast CS+_avs_ vs. CS+_app_ (*p* < .05).

#### Associations with computational learning signals

As a further test of convergence, we re-ran the respective full GAMM (including all design-based factors) with learning signals entered as additional predictors (CS-locked: *V*, α; US-locked: δ+, δ-; each signal as parametric term and difference smooth over *Time* by condition, i.e., *CS Type*). Broadly consistent with the PCA-LMM-based results, this analysis revealed the following significant (via simultaneous confidence bands, *p*s < .05), incremental effects (above experimental conditions): For appetitive cues, pupil dilation covaried positively with expected value (*V*) in a short early cluster (from ca. 0.7 to 0.9 s) after CS onset (descriptively longer, and also present for aversive CS+) and again in a late phase (from 5.6 s onward); yet negatively with associability (α) towards the end of the CS-locked window (from 4.8 s onward). For aversive cues, there was a positive modulation by α in a broad mid-latency cluster (1.2-3.3 s). Positive prediction errors were linked to increased pupil size across most of the post-US window (starting from 0.6 s after US onset) across both valences, whereas negative prediction errors specifically predicted pupil dilation in an earlier cluster (0.7-2.9 s after US onset), which was significant for aversive cues only. For a graphical illustration, see Supplementary Material (Figure S9). Showing a largely convergent pattern of significant associations and their temporal localization, these results thus support the findings from LMM-based analyses.

### 3.6 Out-of-sample validation

Finally, the generalizability of our main findings was assessed using independent, publicly available aversive and appetitive conditioning datasets (Korn et al., 2017; Xia et al., 2025). The temporal component structure of pupillary waveforms generalized remarkably well (Tucker congruence: φs ≥ .99). Functional analyses based on the fixed PCA scoring solution reproduced the main temporal dissociations observed in the primary set of studies, including preferential appetitive response differentiation during the early (Post-CS1) phase (trend-level valence contrast; *p* = .083) and preferential aversive response differentiation around expected outcome onset (*p* < .001). The temporal specificity of design-based effects suggested by the primary analyses was also largely preserved, with deviations from the pattern found in later CS-locked and valence-dependent post-US effects, consistent with differences in timing and paradigm characteristics. Detailed results are reported in the Supplementary Material (Figures S10, S11 and Table S6).

## 4. Discussion

Building on earlier meta-analytic evidence for pupil dilation as a reliable and sensitive marker of Pavlovian conditioning across aversive and appetitive learning (Finke et al., 2021), the present multi-study individual-participant-data reanalysis provides an in-depth examination of its temporal signatures. Based on a large, harmonized dataset from 385 participants (562 conditioning sessions), we demonstrate that the pupillary response during conditioning is not a unitary marker of LC-NE arousal or related neurocognitive processes (Joshi et al., 2016), but rather consists of multiple temporally dissociable components that carry different information about the processes involved in associative learning. Our main conclusions are summarized below. For an overview and tentative integration of our findings see also Table 6.

**Table 6.**
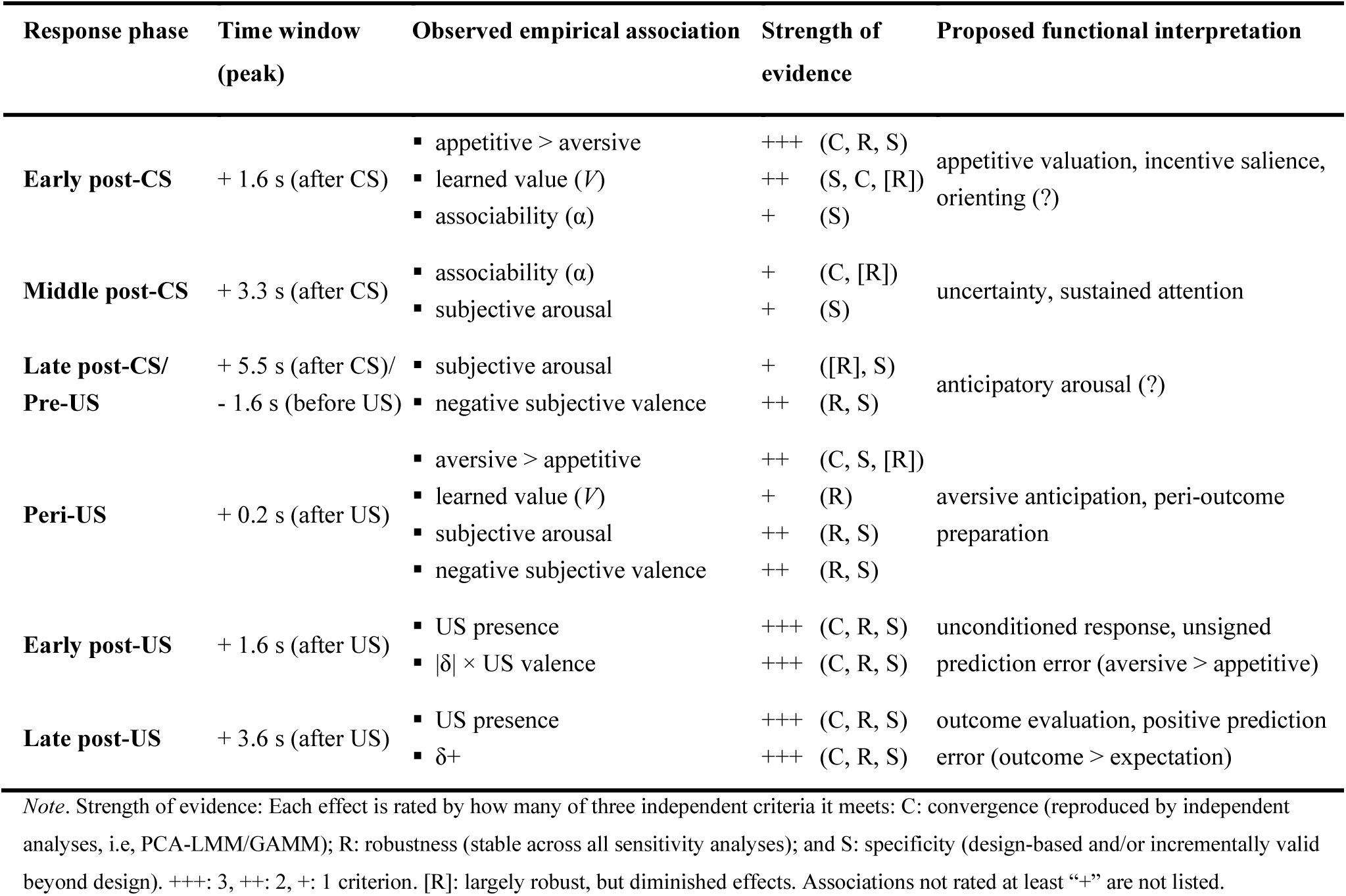
Tentative integration of findings.

### Pupil dilation responses during Pavlovian conditioning are temporally multidimensional and consist of reliably dissociable components

We performed temporal PCA at the trial level to isolate latent components underlying the time course of pupil dilation during both CS-locked and US-locked windows. For each analysis window, PCA revealed a stable component structure that was highly consistent between experimental halves as well as across various contributing samples, indicating that this temporal pattern represents a largely stationary (time-invariant) signal across acquisition (irrespective of amplitude changes), at least given the present timing and design characteristics. The overall structural fit of the chosen PCA solution was further validated by strong correlations of PCA-reconstructed waveforms, empirical grand averages, and GAMM-predicted marginal waveforms (with the notable exception of appetitive difference waves), and was found to be largely robust to the choice of rotation (see Supplementary Material). Importantly, this finding suggests (1) that conditioned pupil dilation reflects a sequence of distinct, yet overlapping processes (or ‘manifolds’; Blini et al., 2024), rather than representing a unitary readout, a pattern also consistent with the established dissociation between early parasympathetic inhibition and later sustained sympathetic activation (Steinhauer & Hakerem, 1992; Finke et al., 2017; Widmann et al., 2018). In addition, our data indicate (2) that these components can be reliably dissociated and (at least in part) mapped onto interpretable aspects of learning.

### Beyond being an unspecific index of arousal, pupil dilation is sensitive to valence-specific processing, depending on timing

Both PCA-based LMM results and GAMM-derived difference waves provided largely converging evidence for valence-specific dissociations between early and later response components: Specifically, pupil dilation in the initial (post-CS) component was found to preferentially track appetitive learning (potentially also involving a graded value signal) and, accordingly, distinguished appetitive CS+ from never reinforced cues (CS-) much earlier than differential responses to aversive CS+ emerged in GAMM predictions. In our data, this effect was particularly robust: it remained (and actually increased) after exclusion of the (only) contingency-instructed sample and was present across remaining studies, including designs with within-subjects variation of valence and constant CS modality (see Figure S5). At least as a trend, it was also present in the data used for external validation. Moreover, both the descriptive pattern of PCA results and GAMM predictions even suggested a linear, valence-dependent gradient in a short window up to 1 s after CS onset (with aversive cues linked to inverse response differentiation, i.e., pupillary constriction). This finding aligns with evidence from instrumental learning tasks suggesting sensitivity of the pupil to nonconscious value judgments (Bijleveld et al., 2011) as well as valence-graded responding in sign-trackers (Schad et al., 2020; for similar pupillary effects of reward prediction in Pavlovian delay conditioning in mice, see Yamada & Toda, 2022). By contrast, responses immediately before/around US onset (in the US-locked analysis) appeared to be more sensitive to aversive conditioning and incrementally predicted subsequent perceptions of unpleasantness as well as arousal beyond the (design-based) value of the cue. Although the underlying mechanisms remain unclear, this pattern may be compatible with a temporal shift from early incentive-/orienting-related processing to later aversive anticipation (see below).

From a meta-science perspective, our results hint at the possibility that larger average effect sizes reported in aversive conditioning research (Finke et al., 2021), or null effects in early (post-CS) as opposed to late (pre-US) windows (Jentsch et al., 2020), may have resulted not only from modality- and intensity-related US effects (given widespread use of electro-tactile stimulation) but also from the early window’s sensitivity to incentive-related value (which might be missed in studies targeting aversive/threat-related learning). Accordingly, our findings may inform future research on appetitive conditioning, which would likely benefit from selecting response intervals based on the observed early-window sensitivity (rather than intervals commonly used in aversive conditioning studies).

### Anticipatory pupil dilation before outcome onset may reflect the shift from uncertainty-related attention (associability) to outcome expectancy

As responses approached (expected) US onset, their functional profile appeared to shift from cue valence toward uncertainty-driven attention (indexed by an association with PH associability, in both LMM- and GAMM-based analyses) and then toward anticipation and outcome expectancy. In line with this, component scores peaking in the late CS window and around US onset predicted self-reported arousal and aligned with expected value (associative strength). Mirroring the appetitive specificity of the Post-CS1 component, the Peri-US component was largely specific to negative valence, as shown both by differential pupillary responses to aversive cues and its strong link to perceived CS unpleasantness. Given both its temporal loading pattern and further analyses based on non-reinforced trials only, our data suggest that both aversive cues and outcomes contributed to this component, reflecting the emergence of a growing anticipatory, threat-related signal, which is further augmented upon US delivery. This assumed dynamic is also consistent with the Peri-US component’s peak around 0.2 s after US onset, as evoked pupil dilation responses are assumed to emerge not much earlier than 500 ms after an event. While this interpretation in terms of a shift in cognitive-affective processing that is differentially reflected in latent pupil-dilation components seems plausible, it is important to note that it is also speculative to some extent. In particular, antecedent learning signals derived from a hybrid RW-PH model did not reliably explain variance beyond the design-based contrast in the CS-locked window (at least not after the initial response phase). However, rather than indicating an absence of learning, this convergence is expected under RW dynamics, where associative strength asymptotically approaches the manipulated contingencies (which points to a limitation of our study design). Given that the component-specific links did not exceed design-based predictors (and may depend on parameter priors, despite evidence for sign stability), it would be premature to consider these pupillary components a selective readout of learning processes until this hypothesis has been directly tested in future research.

### Responses during the window of (expected) US presentation capture independent effects of outcome evaluation and prediction-error-related processes

The clearest evidence for specific computational signals reflected in pupil size emerged in the window after (expected) US onset, where both post-US components tracked the magnitude of expectation violation (i.e., prediction errors). More specifically, in addition to pronounced effects of unconditioned responding to affective outcomes (rewards and aversive stimuli), the earlier component (peaking 1.5-2 s after US onset) indexed both omissions of expected outcomes (Dercksen et al., 2023) and unexpected outcome presentations, with larger effects observed for aversive conditioning. Interestingly, the second post-US component (peaking around 3.5-4 s after US onset) specifically reflected the amplification of responses to affective outcomes by surprise (i.e., the positive prediction error, irrespective of US valence). Prior research has linked pupil dilation to prediction errors mainly in decision-making tasks, for instance, during both decision formation and feedback processing (Colizoli et al., 2018; Colizoli et al., 2026; Van Slooten et al., 2018), with early post-feedback responses sometimes linked to unsigned prediction error signals and later time windows to signed prediction errors. In Pavlovian conditioning, by contrast, evidence for outcome-evoked pupillary prediction-error responses is less consistent (Kim et al., 2025; Stemerding et al., 2022). Our data (i.e., both PCA-LMM and GAMM results) argue in favor of the idea that responses after expected US onset are scaled by general expectation violation (Preuschoff et al., 2011) in a window around 2 s after onset, with later phases potentially indexing prolonged attention to unexpected affective outcomes. Blunted post-US responding in trace (vs. delay) conditioning further suggests that this signal is sensitive to CS-US contiguity, consistent with less certain outcome prediction when cue and outcome are temporally separated. By contrast, the results were not consistent with an account in terms of symmetrical signed prediction-error signaling. Notably, a recent study applying temporal PCA to single-trial pupil data in an associative-memory task isolated a comparable prediction-error-sensitive component (Xue et al., 2026, reviewed preprint). While the emergence of dissociable components sensitive to prediction errors across mnemonic paradigms supports the utility of this approach, the present results do not allow for extrapolating to potential later response phases or assessing whether signed prediction errors (e.g., signed reward prediction errors) may play a role during later stages of outcome processing as well.

### Data-driven decomposition (PCA) and assumption-free modeling of waveforms (GAMM) converge

A central methodological contribution of the present work is the convergence of two complementary analytic strategies, both of which have been applied to pupillometry before (Siegle et al., 2001; Van Rij et al., 2019), but have – to the best of our knowledge – never been implemented, let alone combined, to study pupil dilation in the context of Pavlovian conditioning (for temporal PCA of pupillary data in memory paradigms see Clewett et al., 2020; Xue et al., 2026, reviewed preprint). While temporal PCA reduces the complexity of pupillary waveforms, allowing the derivation of latent components based on a low-dimensional structure, GAMM reconstructs the full waveform without a-priori assumptions about its shape (Wood et al., 2016). Both approaches proved reliable and yielded near-identical marginal waves in all major conditions as well as a high correspondence of difference waves (CS+ vs. CS-), except for the appetitive CS+ contrast (which showed a similar early shape but more sustained dilation in GAMM-based waveforms). The lesser degree of convergence in appetitive conditioning may be related to its smaller signal-to-noise ratio (i.e., smaller differentiation effects), as suggested by additional sensitivity analyses showing relatively lower odd-even reliability for appetitive CS-locked difference waves (*r* = .81, aversive: *r* = .95 [Spearman-Brown corrected]; see Supplementary Material, Figure S8) and thus naturally limited convergent validity with other measures. Overall, dual evidence from two distinct methodological approaches strengthens confidence that the component structure and its association with time-resolved conditioning effects are robust and reflective of genuine empirical relationships (rather than artifacts of the decomposition method or arbitrary analytic choices). In addition, our findings underscore that both methods may be preferable to averaging over broad (and often heterogeneously defined) response windows.

### Limitations and further directions

Our above conclusions are qualified by several noteworthy limitations. First, although all datasets were harmonized and reprocessed using an identical pipeline, the present study relied on a non-preregistered reanalysis of previous (or in part ongoing) studies from a single research group using similar paradigms, which constrains its generalizability. However, this limitation is substantially mitigated by additional, independent out-of-sample validation (added during the peer-review process), showing that the temporal component structure replicated in seven external experiments, and that the functional organization of the components largely generalized as well; this also illustrates that component scores can be projected onto independent data using the present weight matrices, without re-estimating the decomposition (see Data and code availability, below). Extending this framework in future research by systematically addressing the influence of variation in stimulus modality and timing would nonetheless further strengthen the external validity of our findings. Also, given the non-preregistered nature of the present reanalysis, analytical flexibility remains an important limitation. Although we did not conduct an exhaustive multiverse analysis (Lonsdorf et al., 2022), targeted sensitivity and robustness analyses assessed the impact of several core analytical decisions concerning PCA specification (particularly the choice of rotation), inclusion of individual study samples, computational-model parameterization, and the analytical approach used to characterize the pupillary waveform (PCA vs. GAMM). Second, to avoid circularity, learning signals were generated from fixed, plausible starting values, rather than fitted individually. While the observed patterns of associations were quite robust against variation in these parameter choices (in particular regarding the results in the post-US window, which suggested a strong link to prediction-error processing), this might nevertheless have led to reduced overall model fit and a less optimal description of learning trajectories. In a similar vein, collinearity between modeled expected value and experimentally manipulated contingencies (as noted above) restricts inferences about the incremental information conveyed by anticipatory signals, particularly during the CS window. Third, self-report ratings were collected only after acquisition, which constrains their interpretability to global predictive validity between pupillary dynamics and learning outcomes, precluding an in-depth analysis of trial-by-trial convergence with self-reported cognitive-affective processing. Furthermore, all samples comprised healthy, predominantly young adults (with an overall high proportion of women). Therefore, it is unclear whether and how the results generalize to clinical or older populations. Notably, all studies included used timing features (e.g., CS-US onset delay) that are common in conditioning research. However, whether and how divergent designs using very short CS duration (e.g., Roesmann et al., 2022) or rapidly sequenced presentation align with our findings or instead show completely different temporal signatures, is an open question and an interesting avenue for further research.

We performed a series of thorough robustness checks to corroborate our main findings and conclusions. Whereas the early appetitive dissociation and the outcome effects in the post-US window were confirmed as highly stable in these additional analyses (see Supplementary Material), the potential specificity of the Peri-US component to aversive learning was found to be strongly influenced (although not solely driven) by a single study (the only one using explicit contingency instructions). At the same time, the association with self-reported negative cue valence remained robust across samples. Given the fact that instructed fear is likely to potentiate anticipatory responses to the US (presumably peaking shortly before or around its onset), this result is thus consistent with our overall (tentative) interpretation of potential functional correlates underlying this response phase (i.e., expectancy-driven aversive anticipation/preparation). While the less clear-cut pattern of CS vs. US differentiation in the varimax-based sensitivity analysis can be explained by the fact that temporal PCA is blind to the boundary between CS and US intervals, and orthogonal rotation can therefore draw US-related variance into adjacent components (whereas promax permits correlated components and separates this variance more cleanly), findings in the US-locked windows should nonetheless be interpreted with some caution.

On a more general level, our data remain silent as to the specific underlying neurophysiological mechanisms. For instance, rapid effects of cognitive processing on pupil size have been mainly associated with central inhibition of the parasympathetic pathway (Steinhauer et al., 2004), causing relaxation of the sphincter, whereas arousal-induced effects presumably involve direct sympathetic activation of the dilator (Bradley et al., 2008). Similarly, Strauch and colleagues (2022) have more recently proposed a conceptual framework that stipulates distinct types of pupillary responses, distinguished both by their neural underpinnings and (in part) by their temporal characteristics: (a) rapid, transient responses linked to orienting (and presumably subserved by activity in the superior colliculus; see also Wang & Munoz, 2015), (b) relatively delayed, yet more sustained responses indexing alertness (corresponding to activity in the LC-NE system), and (c) more variable effects linked to executive functioning (i.e., the fronto-parietal network). In line with these considerations, we found evidence for similar temporal dynamics (i.e., comparable latencies of component peaks) across both post-CS and post-US response phases. While it is difficult to speculate about the exact neurocognitive underpinnings of the observed valence-dependent dissociation, this framework may also provide one possible account of this pattern. Reward-predictive cues may engage salience- and orienting-related circuitry particularly early, and reward anticipation has been linked to pupil dilation and activity in core salience-network regions (Schneider et al., 2018). In line with this, transient pupil responses may be driven by orienting-related circuitry (Wang & Munoz, 2015). By contrast, aversive differentiation may rely more strongly on anticipatory defense preparation toward expected US onset, consistent with evidence linking aversive anticipation and fear-related pupil dilation to defensive-network regions (Leuchs et al., 2017; Andrzejewski et al., 2019). However, it is important to note that this remains a possible, albeit hypothetical, mechanistic interpretation that is not supported by direct neurophysiological evidence in the present study. Therefore, further research is needed to elucidate if and how distinct sequential stages of pupil dilation during conditioning are linked to specific neurocognitive processes and corresponding patterns of brain activation.

## Conclusion

Taken together, the results of this systematic, large-scale reanalysis of pupillary responses during Pavlovian conditioning are consistent with the idea of a functional dissociation between distinct, early vs. late response components in conditioned pupil dilation. Specifically, early responses seem to be more sensitive to reward-related processing yet less strongly linked to overall levels of arousal. Therefore, our data imply that pupil dilation should not be treated as a unitary marker of associative learning. Rather, selection of response windows should be tailored to the process of interest in future research, rather than assuming identical temporal dynamics in aversive and appetitive conditioning. In line with this, we found initial evidence that these temporally dissociable components may differentially index specific processes such as appetitive value, anticipation, and outcome-related prediction errors, suggesting that decomposing the within-trial temporal signatures of pupil dilation offers a promising, non-invasive approach for future computational modeling applications.

## Author contributions

JBF: conceptualization, methodology, investigation, formal analysis, funding acquisition, visualization, writing (original draft); AMS: investigation, visualization, writing (review and editing); HS: investigation, resources, writing (review and editing); TK: resources, writing (review and editing).

## Author Notes

None of the authors has any potential conflicts of interest to declare. The research was supported (in part) by a grant from the German Research Foundation (DFG) to JBF (grant number FI 2441-4/1).

## Data and code availability

A large part of the individual-participant data analyzed for this study is already openly available in the Open Science Framework (OSF) via repositories of the respective primary studies (https://osf.io/2bnxw; https://osf.io/agjn7; https://osf.io/jbsh4; see also Finke et al., 2023b; Finke et al., 2025; Finke & Klucken, 2025). The temporal PCA weight and loading matrices (CS- and US- locked, promax- and varimax-rotated) are openly available at OSF as well (https://osf.io/m3wnx), allowing projection of component scores onto independent data (a brief implementation guide and easy-to-adapt code example are included). For two ongoing studies as well as one study under review (Schippers et al., 2026), the data cannot be shared publicly prior to publication of the respective primary reports; these data are available from the corresponding author upon reasonable request and will be uploaded on OSF after publication of the primary studies. Additional custom code used in the present analysis is available from the corresponding author upon reasonable request (for academic, non-commercial purposes).

External validation analyses are based on publicly available datasets shared by the originating research group and analyzed, among others, in Korn et al. (2017) and Xia et al. (2025), accessed via Zenodo (CC BY 4.0) and reprocessed using the pipeline described above; the DOIs are listed below. We thank all original authors involved for making these data openly available. Korn et al. (2017): https://doi.org/10.5281/zenodo.1039580; https://doi.org/10.5281/zenodo.5573788; https://doi.org/10.5281/zenodo.1211609; https://doi.org/10.5281/zenodo.3601250; https://doi.org/10.5281/zenodo.7477719; Xia et al. (2025): https://doi.org/10.5281/zenodo.12580446; https://doi.org/10.5281/zenodo.12580463

