## Supplementary Material for "Disentangling the temporal signatures of conditioned pupil dilation: Distinct valence- and prediction-error-related components revealed by mega-analysis"

### Supplementary PCA results

**Dimensionality and component selection.** Figure S1 shows screeplots for the first 15 candidate components based on unrestricted principal component analyses (PCA) in the CS-locked and US-locked windows, respectively. As evident from the shape of the distribution of eigen values, traditional criteria are not well-suited to estimate the dimensionality of pupil traces (i.e., number of components) due to the high autocorrelation inherent in the (10 Hz) data ($\rho \approx .97$). We therefore prioritized consistency/ reproducibility (i.e., Tucker congruence) as well as interpretability.

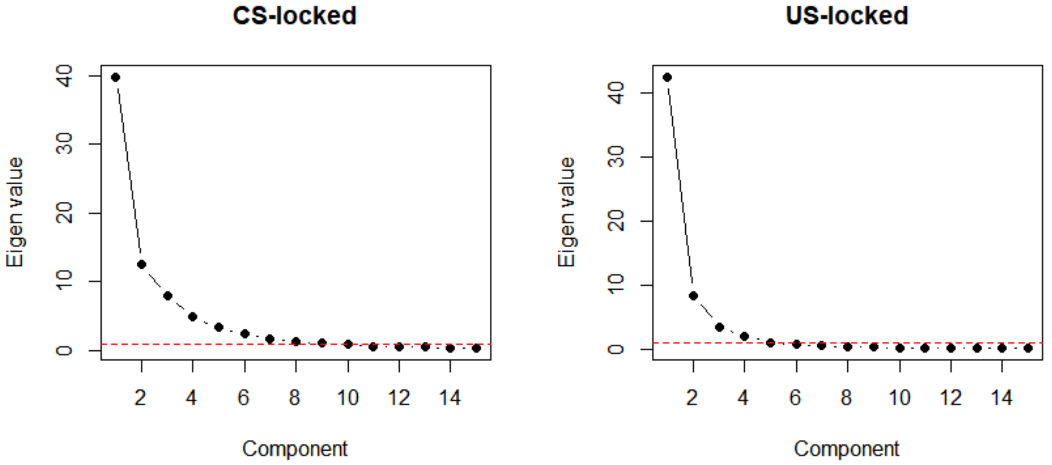


**Supplementary Figure S1.** Screeplots of eigen values of PCAs (first 15 components only) in the CS-locked (left) as well as US-locked window. Red line: Kaiser-Guttman criterion (eigen value > 1).

**PCA solution in the CS-locked window.** After promax rotation, all components during CS presentation were moderately intercorrelated (Post-CS1/Post-CS2: $r = .48$; Post-CS2/Post-CS3: $r = .61$; Post-CS1/Post-CS3: $r = .42$), with the baseline component (Pre-CS) showing notably smaller correlations (all other $r$s $\leq .3$). Note that an alternative solution based on uncorrelated components (using varimax rotation) was highly comparable and led to a very similar overall pattern of results (see Figure S2, below).

All alternative solutions involving a lower number of components explained considerably less variance, whereas solutions with higher number of components showed considerably less stability (Tucker congruence) across studies (see Table S1 for details), indicating that additional components no longer captured a replicable structure.

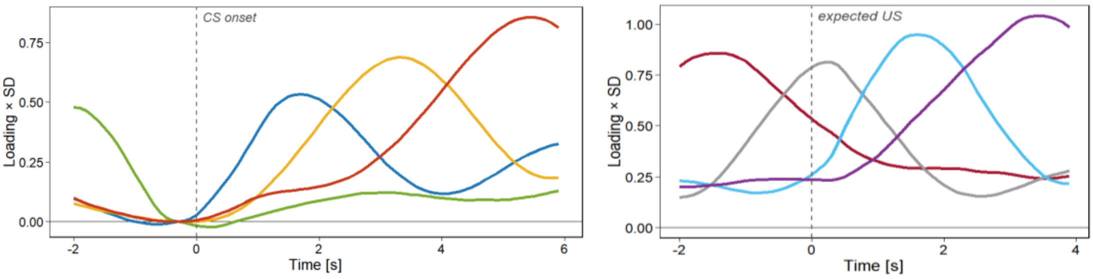


**Supplementary Figure S2.** PCA loading patterns after varimax (instead of promax) rotation in the CS-locked (left) and US-locked (right) windows.

**Supplementary Table S1**. Comparison of stability of PCA solutions in both windows of analysis

| | *CS-locked* | | | | *US-locked* | | | |
|---|---|---|---|---|---|---|---|---|
| **Number of components** | **Median ($\varphi_{min}$)** | **Min. ($\varphi_{min}$)** | **Max. ($\varphi_{min}$)** | ***% variance*** | **Median ($\varphi_{min}$)** | **Min. ($\varphi_{min}$)** | **Max. ($\varphi_{min}$)** | ***% variance*** |
| 1 | 1 | 1 | 1 | 64 | 1 | 1 | 1 | 71 |
| 2 | 1 | 0.99 | 1 | 75 | 0.99 | 0.99 | 1 | 85 |
| 3 | 1 | 0.97 | 1 | 82 | 0.97 | 0.85 | 1 | 91 |
| 4 (selected) | 1 | 0.99 | 1 | 88 | 0.98 | 0.96 | 0.99 | 94 |
| 5 | 0.97 | 0.76 | 0.99 | 91 | 0.82 | 0.46 | 0.98 | 96 |
| 6 | 0.97 | 0.80 | 0.99 | 93 | 0.92 | 0.54 | 0.97 | 97 |

*Note*. $\varphi_{min}$ denotes the smallest Tucker coefficient (i.e., lowest correspondence of components) per run (leave-one-sample-out analysis). Median, minimum and maximum values across all runs are reported.

**PCA solution in the US-locked window.** Adjacent components in the US-locked PCA were strongly correlated (Post-US1/Post-US2: $r$ = .64; Pre-US/Peri-US: $r$ = .62; all other $r$s ≤ .5). Again, the temporal loading pattern of the varimax-solution was highly comparable (see Figure S2), indicating robustness against the choice of rotation.

All alternative solutions involving a higher number of components showed considerably less stability (Tucker congruence) across studies (see Table S1). Similar to the CS-locked analysis, four components thus represented a local stability optimum within the range of stable solutions.

**Within-session stability of PCA solutions: split-half congruence.** Figure S3 illustrates the time invariance of the component structure (for each analysis window), based on the complete dataset. Data were split across the median of the trial variable for each study and compared in terms of Tucker congruence scores, indicating excellent similarity (see main text).

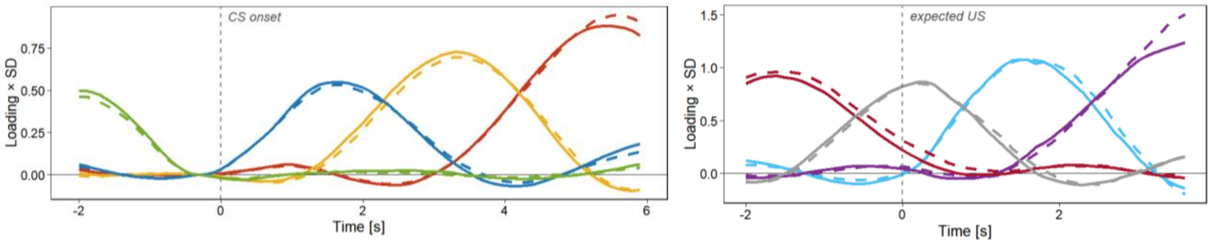


**Supplementary Figure S3.** PCA loading patterns for first (solid lines) vs. second half (dashed lines) of trials in the CS-locked (left) and US-locked (right) windows.

### Supplementary LMM results: further design-based predictors

**CS-locked analysis**. No effects of *CS-US Contiguity* (all $\chi^2$s[1] < 1.72; *p*s ≥ .18) or any *CS-US Contiguity* × *CS Type* interactions (all $\chi^2$s[3] < 4.29, *p*s ≥ .23) emerged for any CS-locked component. The effect of *CS Trial* was significant for the early and middle components (Post-CS1: $\chi^2$[1] = 6.22; *p* = .013; *b* = -0.003, *SE* = 0.001, *t*[23300] = -2.50, *p* = .013; Post-CS2: $\chi^2$[1] = 5.49; *p* = .019; *b* = -0.003, *SE* = 0.001, *t*[23480] = -2.34, *p* = .019). However, this trend across acquisition was completely unrelated to differential responding (i.e., not dependent on *CS Type*; $\chi^2$s[2] ≤ 2.3; *p*s ≥ .31). For the late component (Post-CS3), there was no indication of any change across trials at all ($\chi^2$[2] = 0.31; *p* = .854; interaction: $\chi^2$[1] = 0.67, *p* = .410).

**US-locked analysis**. *CS-US Contiguity* impacted significantly on scores of the earlier, post-CS components (Pre-US: $\chi^2$[1] = 48.37, *p* < .001; Peri-US: $\chi^2$[1] = 10.05, *p* < .001), but this did not interact with *CS Type* (Pre-US: $\chi^2$[2] = 0.49; *p* = .785; Peri-US: $\chi^2$[2] = 1.62; *p* = .445), indicating no influence on differential conditioning. Effects of *CS Trial* on these two components were not significant, either, and neither were *CS Trial* × *CS Type* interactions ($\chi^2$[3]s < 3.98; *p*s > .26). The *CS-US Contiguity* × *US Presence* interaction was significant for both late-peaking components (all $\chi^2$[1]s > 57, *p*s < .001), indicating substantially blunted responding to outcomes in trace conditioning (Post-US1: *b* = -0.491, *SE* = 0.035, *t*[22680] = -13.98, *p* < .001; Post-US2: *b* = -0.277, *SE* = 0.036, *t*[22700] = -7.56, *p* < .001). The effect of *CS Trial* was significant for the Post-US1 component ($\chi^2$[1] = 31.4; *p* < .001; *b* = -0.006, *SE* = 0.001, *t*[22690] = -5.61, *p* < .001; Post-US2: all $\chi^2$s < 0.3, *p*s > .80), but the interaction was not (*CS Trial* × *US Presence,* Post-US1: $\chi^2$[1] = 2.48; *p* = .115; *b* = -0.004, *SE* = 0.002, *t*[22680] = -1.58, *p* = .115).

### Robustness check 1: PCA solution based on orthogonal rotation (‘varimax’)

To provide further evidence for the robustness of the extracted component structure as well as its differential associations with learning-related variables, we additionally conducted PCA using varimax rotation (instead of promax) within each window of analysis (CS-locked, US-locked), yielding rotated yet uncorrelated components. The resulting loading patterns are illustrated in Figure S2, showing high resemblance with the promax-based solution (with the initial Post-CS component being slightly less temporally distinct). All major analytical steps were then re-run in order to corroborate our findings. The results (see below) clearly indicate that the core conclusions, and in particular the full pattern of CS-locked effects, did not depend on this analytical choice. Rather, all primary findings remain robust with uncorrelated components, i.e., under orthogonal (rather than oblique) rotation. The oblique (promax) solution, however, achieved a somewhat cleaner separation of responses, especially around US onset.

**Tucker congruence.** Split-half congruence of the PCA solution was near-perfect for both windows of analysis (all Tucker φs ≥ .98). Congruence across different studies (based on an iterative leave-one-sample-out approach) was likewise excellent for both windows (φs ≥ .96).

**Linear mixed-effects models (LMMs).** Tables S2 and S3 report the results of LMMs fitted to scores of each (varimax-based) component, including all core design-based predictors. The pattern of statistically significant results in the CS-locked window was equivalent to the outcome of the main analysis, apart from a minor, trend-level difference between the CS+ and CS- that emerged for the Pre-CS component (baseline) and presumably resulted from a small leakage effect (due to the less clean temporal separation; see Figure S2). While the overall pattern of primary CS- vs. US-related effects was also stable in the US-locked window (see Table S3), variance linked to US valence appeared more broadly distributed across components (consistent with the less simple loading structure). Crucially, however, the differential valence contrast for Peri-US scores remained significant ($CS+_{avs} > CS+_{app}$: $p < .001$). Taken together, the results indicate that the varimax rotation, despite its overall similarity, was not as successful as the promax rotation in decomposing responses in the window around US onset in a way that mapped cleanly onto the CS/US distinction.

**Association with computational learning signals.** As shown in Figure S4 (A), the pattern of correlations with learning signals in the CS-locked window was highly similar to the main analysis. Only the Pre-US component showed a markedly different profile, with novel, divergent associations emerging with $V$ (positive) and α (negative), similar to the Peri-US component. Notably, this may be directly related to the somewhat less clear dissociation between CS-evoked and US-/outcome-evoked responses in the varimax-based PCA solution (see above). For both post-US components, an almost identical pattern of modulation by unsigned prediction errors (as reported in the main analysis) was found (see Figure S4, C)

**Association with self-report learning indices.** As shown in Figure S4 (B), the pattern of correlations with self-report indices was largely equivalent to the outcome of the main analysis. Again, only the Pre-US component showed a different profile. Moreover, arousal levels attributed to the CS were less reliably predicted by the middle (Post-CS2/Post-CS3) components.

**Supplementary Table S2.** Model parameters of primary linear mixed-effects models predicting (varimax-based) component scores in the CS-locked window

| | **Component** (incl. peak latency) | | | | | | | | | | | | | | | | | | | |
|---|---|---|---|---|---|---|---|---|---|---|---|---|---|---|---|---|---|---|---|---|
| | ***Pre-CS (~ -2.0 s)*** | | | | | ***Post-CS1 (~ +1.7 s)*** | | | | | ***Post-CS2 (~ +3.3 s)*** | | | | | ***Post-CS3 (~ +5.5 s)*** | | | | |
| **Model (predictors)** | *b* | *SE* | df | *t* | *p* | *b* | *SE* | df | *t* | *p* | *B* | *SE* | df | *t* | *p* | *b* | *SE* | df | *t* | *p* |
| Intercept | 0.001 | 0.010 | 5.66 | 0.15 | .883 | 0.006 | 0.025 | 6.09 | 0.23 | .826 | 0.009 | 0.024 | 5.75 | 0.36 | .730 | -0.024 | 0.066 | 5.94 | -0.35 | .732 |
| *CS Type* | | | | | | | | | | | | | | | | | | | | |
| CS+ (vs. CS-) | -0.024 | 0.014 | 15330 | -1.80 | .072 | -0.005 | 0.013 | 23300 | -0.41 | .681 | 0.043 | 0.013 | 23280 | 3.26 | **.001** | 0.035 | 0.013 | 23470 | 2.66 | **.008** |
| Appetitive (vs. aversive) | 0.030 | 0.019 | 184.3 | 1.61 | .109 | 0.090 | 0.020 | 6186 | 4.58 | <**.001** | -0.008 | 0.020 | 5915 | -0.40 | .692 | -0.029 | 0.020 | 22430 | -1.49 | .137 |
| *Variance Components* | $\sigma^2$ | *SD* | | | | $\sigma^2$ | *SD* | | | | $\sigma^2$ | *SD* | | | | $\sigma^2$ | *SD* | | | |
| Sample | 0.000 | 0.015 | | | | 0.003 | 0.057 | | | | 0.003 | 0.058 | | | | 0.029 | 0.171 | | | |
| Subject | 0.003 | 0.051 | | | | 0.023 | 0.152 | | | | 0.015 | 0.122 | | | | 0.024 | 0.156 | | | |
| Subject:Session | 0.001 | 0.027 | | | | 0.009 | 0.096 | | | | 0.009 | 0.096 | | | | 0.012 | 0.109 | | | |
| Residual | 1.017 | 1.009 | | | | 0.981 | 0.990 | | | | 0.973 | 0.987 | | | | 0.968 | 0.984 | | | |

*Note*. Number of samples: 7; number of subjects: 383; number of sessions: 562; number of trials: 23980. *b*: estimate (component score); *SE*: standard error. *p* values significant at the $p < .05$ level are shown in bold.

.

**Supplementary Table S3.** Model parameters of primary linear mixed-effects models predicting (varimax-based) component scores in the US-locked window

| | **Component** (incl. peak latency) | | | | | | | | | | | | | | | | | | | |
|---|---|---|---|---|---|---|---|---|---|---|---|---|---|---|---|---|---|---|---|---|
| | ***Pre-US (~ -1.4 s)*** | | | | | ***Peri-US (~ +0.3s)*** | | | | | ***Post-US1 (~ +1.6 s)*** | | | | | ***Post-US2 (~ +3.4 s)*** | | | | |
| **Model (predictors)** | *b* | *SE* | df | *t* | *p* | *b* | *SE* | df | *t* | *p* | *b* | *SE* | df | *t* | *p* | *b* | *SE* | df | *t* | *p* |
| Intercept | -0.028 | 0.052 | 5.89 | -0.54 | .610 | 0.028 | 0.062 | 5.91 | 0.45 | .671 | -0.007 | 0.123 | 6.04 | -0.06 | .954 | 0.047 | 0.072 | 5.77 | 0.66 | .537 |
| *CS Type* | | | | | | | | | | | | | | | | | | | | |
| CS+ (vs. CS-) | 0.045 | 0.018 | 22640 | 2.54 | **.011** | 0.041 | 0.018 | 22700 | 2.28 | **.023** | -0.007 | 0.016 | 22630 | -0.45 | .651 | -0.010 | 0.017 | 22650 | -0.60 | .549 |
| Appetitive (vs. aversive) | 0.006 | 0.024 | 20670 | 0.26 | .798 | -0.135 | 0.024 | 21760 | -5.58 | <**.001** | -0.094 | 0.022 | 22610 | -4.21 | <**.001** | 0.109 | 0.023 | 22210 | 4.71 | <**.001** |
| *US Presence* | | | | | | | | | | | | | | | | | | | | |
| US+ (vs. US-) | -0.279 | 0.018 | 22660 | -10.05 | <**.001** | -0.093 | 0.018 | 22710 | -5.19 | <**.001** | 0.462 | 0.017 | 22640 | 27.96 | <**.001** | 0.356 | 0.017 | 22660 | 20.76 | <**.001** |
| Appetitive (vs. aversive) | 0.021 | 0.035 | 22660 | 0.60 | .549 | 0.121 | 0.035 | 22710 | 3.42 | <**.001** | 0.004 | 0.033 | 22640 | 0.13 | .896 | -0.114 | 0.034 | 22660 | -3.36 | <**.001** |
| *Variance Components* | $\sigma^2$ | *SD* | | | | $\sigma^2$ | *SD* | | | | $\sigma^2$ | *SD* | | | | $\sigma^2$ | *SD* | | | |
| Sample | 0.018 | 0.134 | | | | 0.026 | 0.163 | | | | 0.087 | 0.295 | | | | 0.035 | 0.187 | | | |
| Subject | 0.011 | 0.106 | | | | 0.014 | 0.117 | | | | 0.042 | 0.207 | | | | 0.010 | 0.100 | | | |
| Subject:Session | 0.040 | 0.201 | | | | 0.010 | 0.099 | | | | 0.040 | 0.200 | | | | 0.055 | 0.235 | | | |
| Residual | 0.946 | 0.973 | | | | 0.949 | 0.974 | | | | 0.809 | 0.900 | | | | 0.874 | 0.935 | | | |

*Note.* Number of samples: 7; number of subjects: 385; number of sessions: 557; number of trials: 23181. *b*: estimate (component score); *SE*: standard error. *p* values significant at the $p < .05$ level are shown in bold.

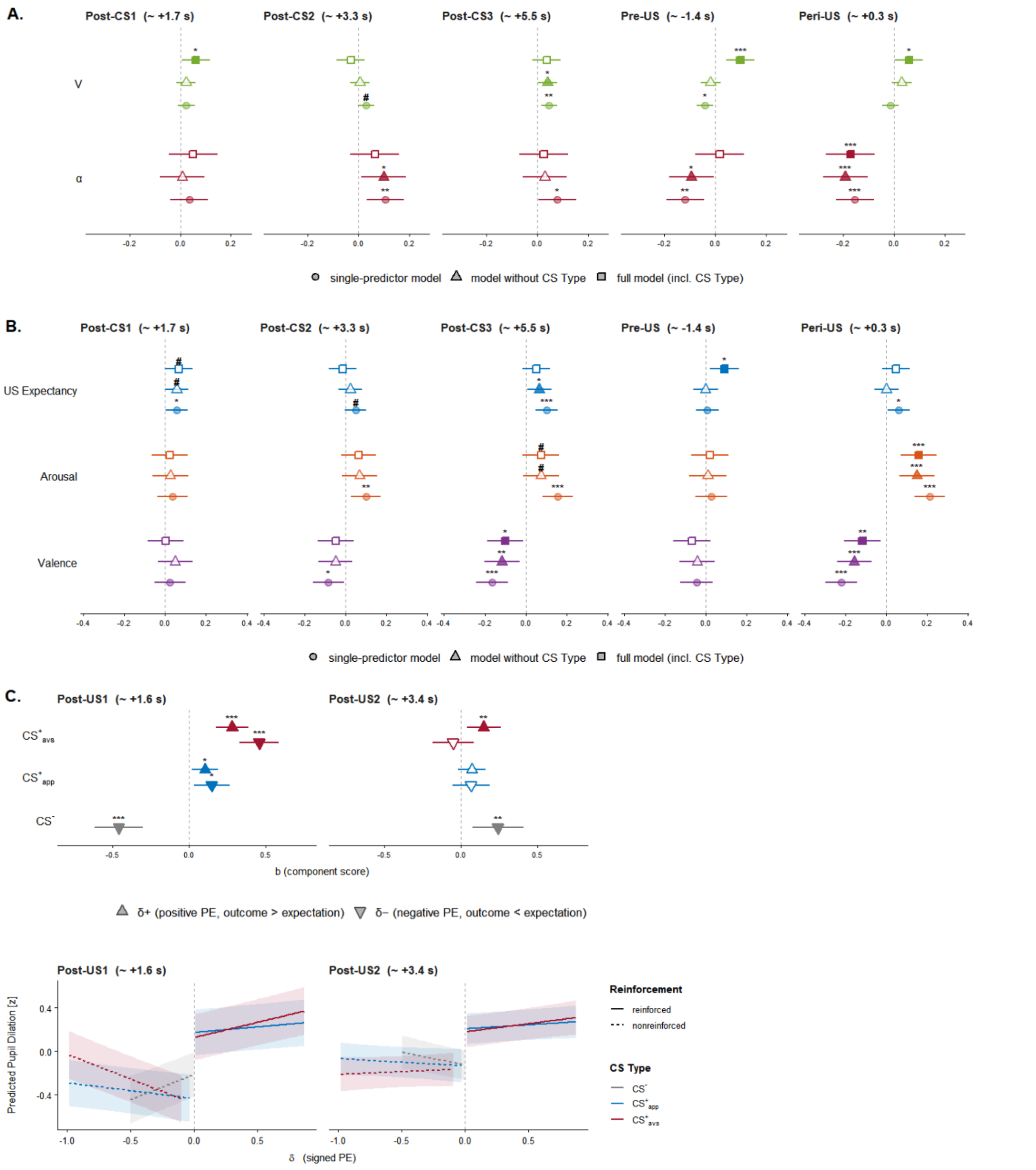


**Supplementary Figure S4**. Associations (based on LMMs) of A. antecedent learning signals (value, *V*; associability, α) and B. self-report learning indices (*US Expectancy*, *Arousal* and *Valence* ratings) with conditioned pupil dilation across varimax-rotated PCA components (before/around US onset).

*Note.* X-axis: *b* estimate (component score). Squares: estimates based on full models (A. model 3 [cf. Table 4]: *score* ~ *CS_Type* + *V* + α + random effects; note that the coefficient shown for Post-CS1/*V* is based on the pooled effect for the CS+, due to the significant *V* × *CS Type* interaction; B. model 6: *score* ~ *CS_Type* + *US_Expectancy* + *Arousal* + *Valence* + random effects); triangles: estimates based on models without design-based predictors (A. model 2: *score* ~ *V* + α + random effects; B. model 5: *score* ~ *US_Expectancy* + *Arousal* + *Valence* + random effects); circles: simple mixed-model regression estimates for each predictor. Error bars indicate 95% CI.

C. Differential associations (by levels of *CS Type*) with unsigned prediction errors (PE) for response components after (expected) US onset. Grey line: simple PE effect (without interaction).

*Note.* $CS+_{avs}$: aversive conditioned stimulus; $CS+_{app}$: appetitive conditioned stimulus; CS-: never reinforced control stimulus. Error bars indicate 95% CI.

#### Robustness check 2: Distribution of effects across primary studies

To rule out that the pattern of results, especially concerning the valence-specific effects, was driven by individual samples/studies, we performed a further sensitivity analysis and fitted all primary linear mixed-effects models separately for each study (i.e., by Sample ID). Figure S5 shows the resulting distribution of contrast coefficients (CS+ vs- CS-) per level of *US Valence* for each component, also including the US contrast (present vs. absent) for components in the post-US window. As evident from this illustration, the pattern was largely consistent across studies, except for Sample ID 1, which was the only study showing a positive effect for the aversive CS+ in the early (Post-CS1) component (as well as higher aversive CS contrasts overall). Notably, this was also the only study which applied a full contingency instruction (i.e., information about the correct contingencies before acquisition).

To account for this potential source of bias, we repeated all analyses after exclusion of this sample, which in fact revealed even stronger valence-specific effects in the Post-CS1 component (see Figure S5). Almost all effects (as reported in Tables 2 & 3 of the main text) remained significant and virtually unchanged, except for the CS contrasts in the Post-CS3 and Pre-US components which were strongly diminished, especially with aversive cues. Importantly, the pattern of aversive-specific differential responding observed for the Peri-US component was robust as well (aversive: $p = .030$; appetitive: $p = .372$), and its associations (in the respective full model) with self-report learning indices remained significant, i.e., the negative correlation with subjective cue valence ($p = .008$) and positive correlation with subjective arousal ($p = .036$), respectively. Notably, associations with computational learning signals were also reliable, including Post-CS1/$V$ and $\alpha$ as well as prediction-error modulation of Post-US1 and Post-US2 scores (all $ps < .025$), except for Post-CS2/$\alpha$ which was significant as single predictor, but not in the joint model. Taken together, this outcome supports our main conclusions.

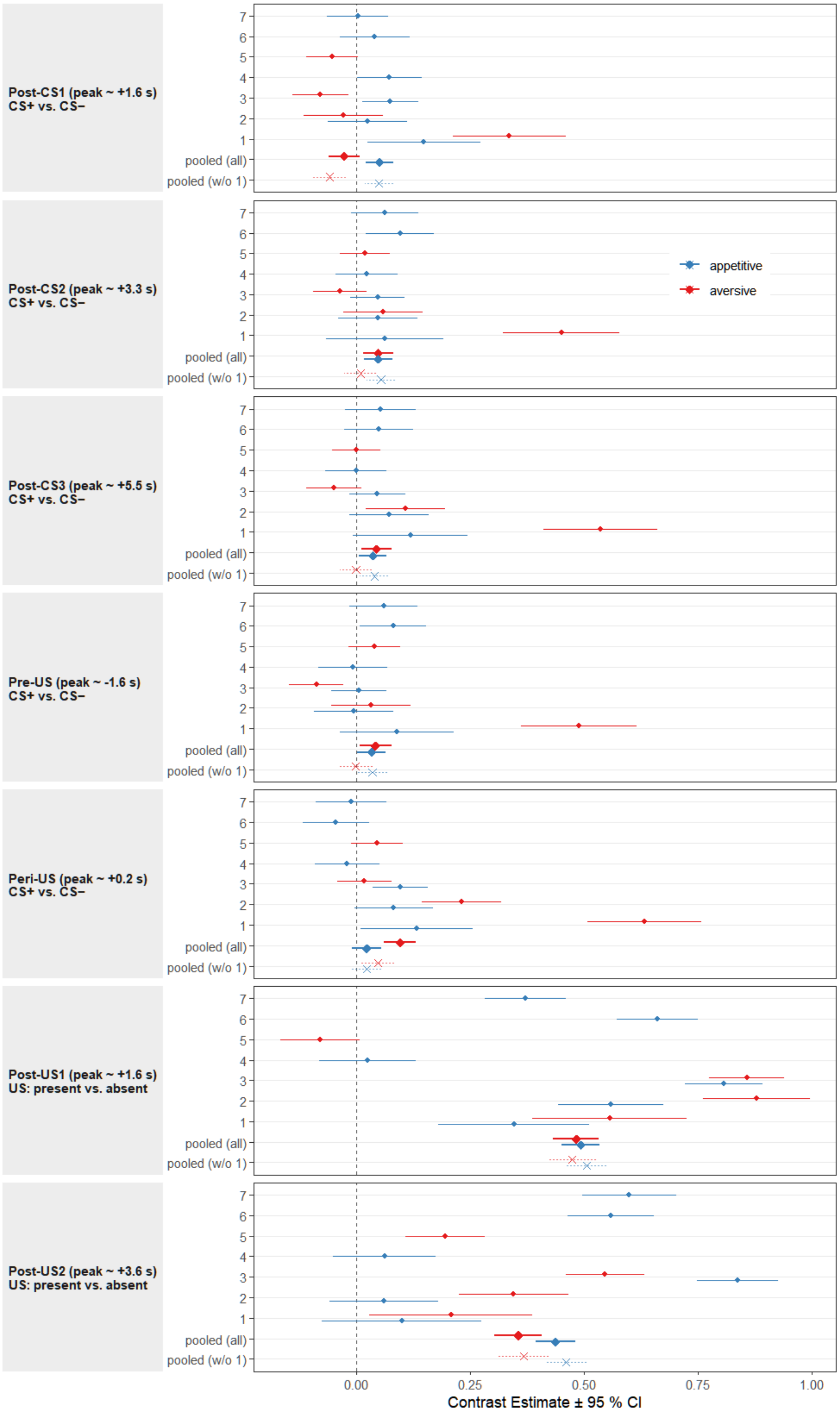


**Supplementary Figure S5.** Distribution of effects (main contrasts) across samples (primary studies) for each component.

### Robustness check 3: Parameter choice for computational learning model

Because the initial values of the RW-PH learning model ($V_0$, $\alpha_0$, $\gamma$) were fixed a priori to avoid inferential circularity (i.e., comparison with the same data used for fitting), rather that estimated from individual model fits, we performed a sensitivity analysis in which all learning signals were recomputed across a grid of plausible starting values ($V_0$: [0, 0.125, 0.25, 0.375, 0.5]; $\alpha_0$: [0.4, 0.5, 0.6, 0.7, 0.8]; $\gamma$: [0.02, 0.05, 0.1, 0.2, 0.3]). At each combination of fixed parameter values, the linear mixed-effects model for each component of the pupil response (i.e., best-fitting model as reported in the main text) was refitted. To facilitate comparisons, standardized coefficients were calculated for each effect and are reported both in aggregated form (i.e., range and median; see Table S4) and in full (see Figure S6). Direction and magnitude of all essential learning effects in the post-US window (i.e., Post-US1: $\delta+ \times$ *CS Type*, $\delta- \times$ *CS Type*; Post-US2: $\delta+$ vs. $\delta-$) were preserved throughout estimations (sign stability $\approx 1$; see Table S4). Moreover, most associations observed for CS-evoked responses were highly robust as well, in particular the Post-CS2/$\alpha$ and Peri-US/$V$ relationships. Though consistently sign-stable, the $V \times$ *CS Type* interaction found for the Post-CS1 component was more dependent on initial values, being significant in only 62% of models. Taken together, the results indicate that the overall pattern of associations reported, including major findings, does not depend solely on the choice of fixed values.

**Supplementary Table S4.** Distribution of standardized regression coefficients for the relationship between learning signals and pupil dilation responses, estimated at different combinations of fixed parameter values ($V_0$, $\alpha_0$, $\gamma$) for each PCA component

| Component | *Predictor* | $\beta_{ref}$ | $\beta_{med}$ | $\beta_{min}$ | $\beta_{max}$ | *Prop. stable sign* | *Prop. sig.* |
|---|---|---|---|---|---|---|---|
| Post-CS1 | $V$ [CS+] | 0.031 | 0.026 | 0.010 | 0.031 | 1 | 0.62 |
| | $V$ [CS-] | -0.087 | -0.113 | -0.395 | -0.017 | 1 | 0.44 |
| | $\alpha$ | 0.027 | 0.022 | -0.001 | 0.036 | 0.99 | 0.52 |
| Post-CS2 | $V$ | 0.007 | 0.007 | -0.001 | 0.011 | 0.95 | 0 |
| | $\alpha$ | 0.018 | 0.019 | 0.011 | 0.023 | 1 | 0.84 |
| Post-CS3 | $V$ | 0.016 | 0.013 | 0.002 | 0.023 | 1 | 0.28 |
| | $\alpha$ | 0.009 | 0.011 | -0.003 | 0.017 | 0.97 | 0.13 |
| Pre-US | $V$ | 0.009 | 0.012 | 0.006 | 0.020 | 1 | 0.11 |
| | $\alpha$ | 0.006 | 0.007 | -0.004 | 0.011 | 0.90 | 0 |
| Peri-US | $V$ | 0.024 | 0.023 | 0.017 | 0.025 | 1 | 0.96 |
| | $\alpha$ | -0.003 | 0.001 | -0.005 | 0.008 | 0.58 | 0 |
| Post-US1 | $\delta-$ [$CS+_{app}$] | -0.033 | -0.038 | -0.049 | -0.020 | 1 | 0.89 |
| | $\delta-$ [$CS+_{avs}$] | -0.086 | -0.086 | -0.119 | -0.054 | 1 | 1 |
| | $\delta-$ [CS-] | 0.087 | 0.104 | 0.028 | 0.415 | 1 | 0.78 |
| | $\delta+$ [$CS+_{app}$] | 0.027 | 0.038 | 0.015 | 0.055 | 1 | 0.93 |
| | $\delta+$ [$CS+_{avs}$] | 0.078 | 0.080 | 0.039 | 0.120 | 1 | 1 |
| Post-US2 | $\delta-$ | -0.017 | -0.014 | -0.024 | 0.003 | 0.99 | 0 |
| | $\delta+$ | 0.029 | 0.035 | 0.018 | 0.049 | 1 | 1 |

*Note*. Ref, reference model (based on initial values as reported in the main text); med/min/max, median/minimum/maximum across all estimations; prop. stable sign, proportion of models with identical sign (+ vs. -) of the respective slope; prop. sig., proportion of models with significant slope (at $p < .05$). For the Post-CS1 component, the interaction of $V$ with the *CS Type* contrast was modeled. For the Post-US1 component, $\delta$ slopes are likewise shown separately for each level of *CS Type* (excluding the reference level).

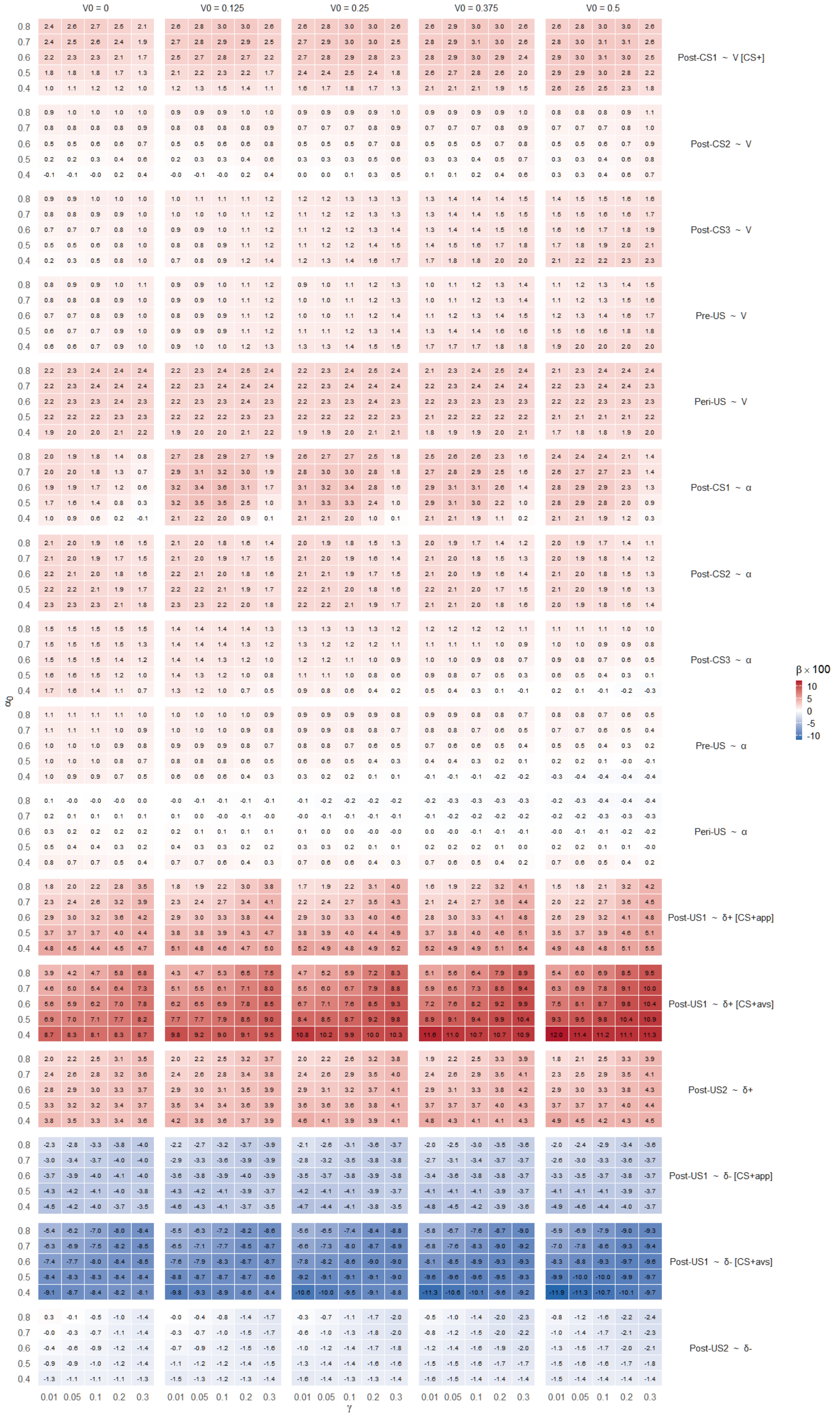


**Supplementary Figure S6.** Sensitivity of the learning-signal associations to fixed RW-PH starting values: Standardized regression coefficients (shown as β×100 for clarity) estimated at different combinations of fixed parameter values ($V_0$, $\alpha_0$, γ) for each component of the pupil response (for model specifications, see main text as well as Table S5). For the Post-US1 component, δ slopes are shown separately for both types of CS+ (appetitive vs. aversive).

### Generalized additive mixed models: Detailed results

Final parameters of the generalized additive mixed models (GAMMs) fitted to pupil dilation responses in the CS-locked and US-locked windows, are reported in Table S5. Figure S7 shows a comparison of GAMM-predicted waveforms with empirical and PCA-based grand averages, illustrating convergence of model fits. In addition, we performed split-half reliability analyses by estimating two separate models for each window, splitting trials (within each CS type) into odd- and even-numbered trials. The results are shown in Figure S8, demonstrating high internal consistency of the GAMM fit.

**Supplementary Table S5.** Model parameters of generalized additive mixed models fitted to pupil dilation responses

| | ***CS-locked*** | | | | | ***US-locked*** | | | | |
|---|---|---|---|---|---|---|---|---|---|---|
| **Model terms** | *b* | *SE* | | *t* | *p* | *b* | *SE* | | *t* | *p* |
| *Parametric coefficients* | | | | | | | | | | |
| Intercept [= CS-] | 0.130 | 0.025 | | 5.23 | <**.001** | 0.298 | 0.037 | | 8.11 | <**.001** |
| *Condition* | | | | | | | | | | |
| CS+$_{avs}$ | 0.016 | 0.004 | | 3.79 | <**.001** | 0.160 | 0.008 | | 21.05 | <**.001** |
| CS+$_{avs}$ \| US+ | - | - | | - | - | -0.041 | 0.008 | | -5.08 | **.003** |
| CS+$_{app}$ | 0.002 | 0.005 | | 0.29 | .769 | 0.103 | 0.010 | | 10.86 | <**.001** |
| CS+$_{app}$ \| US+ | - | - | | - | - | 0.226 | 0.009 | | 23.97 | <**.001** |
| CS-US Contiguity \| trace | 0.000 | 0.011 | | 0.02 | .986 | -0.047 | 0.018 | | -2.65 | **.008** |
| Startle Probe \| present | 0.037 | 0.007 | | 4.91 | <**.001** | 0.128 | 0.010 | | 12.96 | <**.001** |
| | edf | Ref. df | *SD* | *F* | *p* | edf | Ref. df | *SD* | *F* | *p* |
| *Smooth terms* | | | | | | | | | | |
| Time [= CS-] | 23.76 | 23.94 | 0.225 | 71.22 | <**.001** | 23.51 | 23.87 | 0.247 | 33.95 | <**.001** |
| Time \| CS+$_{avs}$ | 18.31 | 21.77 | 0.063 | 6.30 | <**.001** | 17.01 | 20.85 | 0.108 | 3.43 | <**.001** |
| Time \| CS+$_{avs}$ \| US+ | - | - | - | - | - | 23.53 | 23.97 | 0.453 | 143.94 | <**.001** |
| Time \| CS+$_{app}$ | 20.55 | 23.05 | 0.079 | 7.79 | <**.001** | 19.25 | 22.42 | 0.131 | 8.81 | <**.001** |
| Time \| CS+$_{app}$ \| US+ | - | - | - | - | - | 23.37 | 23.95 | 0.336 | 153.33 | <**.001** |
| CS Trial | 6.80 | 7.91 | 0.013 | 4.28 | <**.001** | 6.64 | 7.77 | 0.016 | 3.73 | <**.001** |
| Time \| trace | 11.90 | 13.44 | 0.056 | 5.54 | <**.001** | 13.78 | 13.99 | 0.295 | 63.67 | <**.001** |
| Time \| Startle present | 13.66 | 13.97 | 0.169 | 63.45 | <**.001** | 13.82 | 13.99 | 0.337 | 75.44 | <**.001** |
| *Variance Components* | | | | | | | | | | |
| Sample | 5.48 | 7 | 0.061 | 17.52 | <**.001** | 5.31 | 6 | 0.088 | 19.45 | <**.001** |
| Subject:Time$_1$ | 2614.5 | 3447 | 0.108 | 5.19 | <**.001** | 2960.3 | 3464 | 0.188 | 11.23 | <**.001** |
| Subject:Time$_2$ | | | 0.077 | | | | | 0.135 | | |
| Subject:Session | 305.81 | 561 | 0.100 | 1.53 | <**.001** | 333.0 | 555 | 0.177 | 2.31 | <**.001** |
| Residual | | | 0.548 | | | | | 0.624 | | |
| Deviance explained: | 6.94% | | | | | 11.6% | | | | |

*Note*. Number of sampling points, CS-locked: 1935364; US-locked: 1411077; number of samples: 7; number of subjects, CS-locked: 383, US-locked: 385; number of sessions, CS-locked: 562, US-locked: 557; number of trials, CS-locked: 23980, US-locked: 23181. *b*: estimate (pupil dilation); *SE*: standard error. *p* values significant at the $p < .05$ level are shown in bold. Deviance explained reflects variance explained by fixed effects (most systematic variance is captured by random smooths).

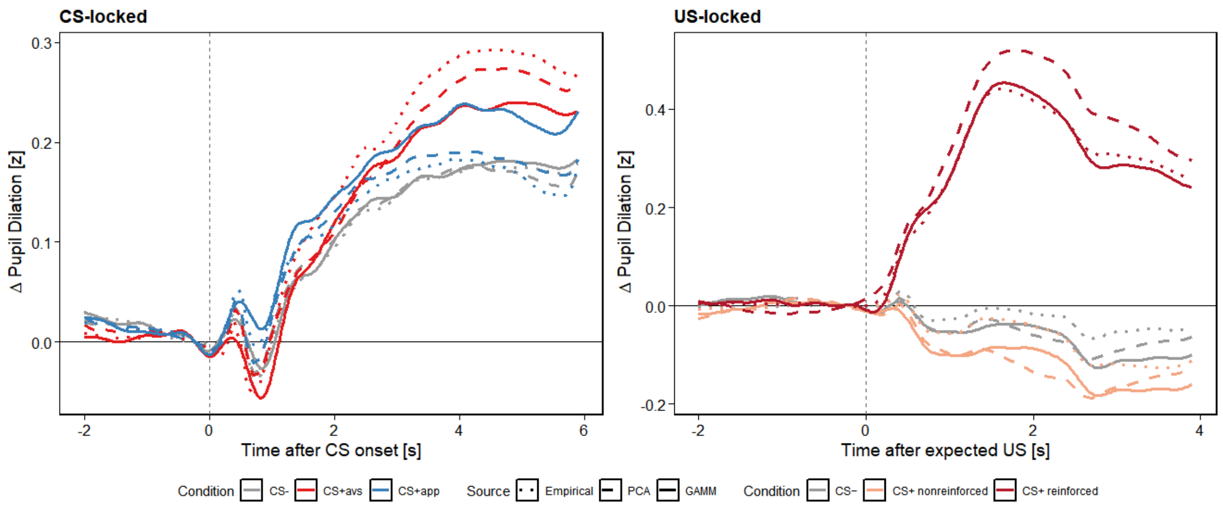


**Supplementary Figure S7.** Overlay of average pupillary waveforms in each major condition (CS-locked, left: appetitive CS+, aversive CS+, CS-; US-locked, right: reinforced CS+ trials, non-reinforced CS+ trials, CS- trials), as indicated by different methods: (1) empirical grand average, (2) mean waveforms reconstructed from PCA, (3) GAMM-predicted marginal waveforms (at reference levels of additional parametric and non-parametric terms). Note that, for reasons of clarity and comparability, waveforms of responses in the US-locked window were re-referenced to the level immediately before US onset.

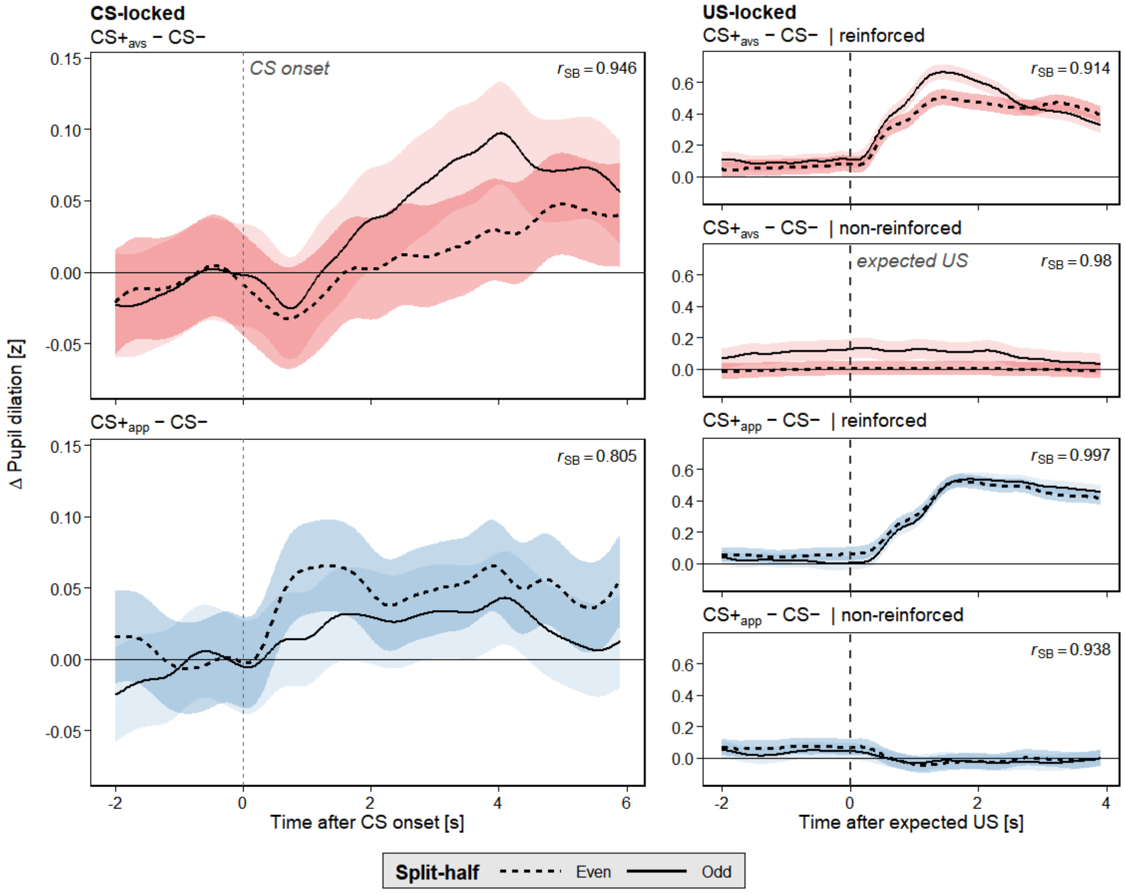


**Supplementary Figure S8.** Comparison (split-half reliability) of GAMM-predicted difference waves (CS+ vs. CS-), separately fitted on either odd- or even-numbered trials. $r$ values shown (indexing shape consistency) are Spearman-Brown corrected.

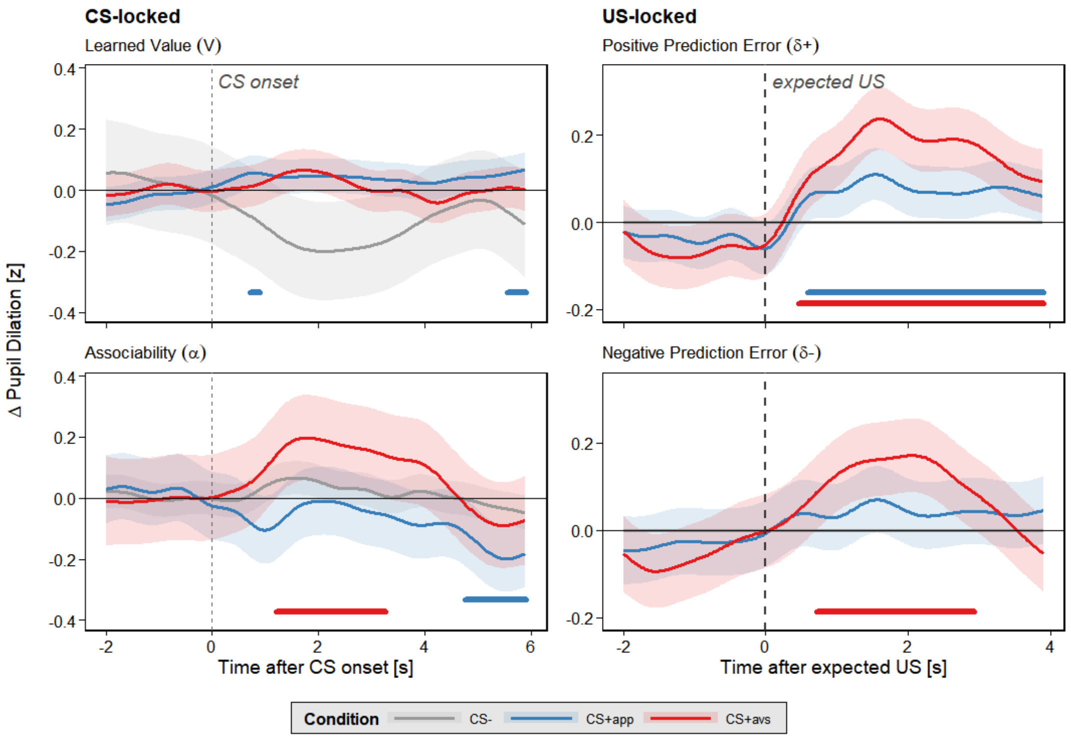


**Supplementary Figure S9.** Associations of computational learning signals (incremental effects above design-based factors) with GAMM-based waveforms in both windows of analysis. Shaded bands: 95% CI. Colored bars at the bottom indicate significant differences from zero, i.e., effects of learning variables in each condition, based on simultaneous 95% CIs ($p < .05$). The difference between the 5% and 95% quantile is used as unit for each predictor. See main text for further explanation.

### Out-of-sample validation

**Method.** To provide additional evidence for the generalizability of our findings, we performed an out-of-sample validation test using publicly available raw pupillometric data from seven conditioning experiments conducted by an independent research group (Korn et al., 2017; Xia et al., 2025). The external set of studies comprised five aversive-conditioning datasets (using electric shock as US; cumulative sample size after exclusions: $n$ = 87) and two appetitive-conditioning datasets (fruit juice; $n$ = 71) and included several CS modalities. Reinforcement rate (CS+ trials) in all studies was 50% (thus similar but slightly lower than the average in the experimental protocols included in our primary analysis). Raw data were harmonized and preprocessed according to the pipeline outlined in the main text (including trial- and participant-level exclusion criteria).

(1) *Structural replication* was assessed by independently estimating four-component covariance PCAs equivalent to the main analysis (separately for CS-locked as well as US-locked data, using promax rotation and the weighting procedure outlined in the main text). The resulting component structure was matched to the corresponding components of the primary solution, and correspondence was quantified using Tucker congruence. Because the timing of outcome delivery differed substantially between the primary and external datasets, with expected or actual US onset occurring within the predefined CS-locked window (-2 to 6 s relative to CS onset) in six of seven external experiments (US onset: +3.5 to

+5 s; +6 s in the remaining experiment), only nonreinforced trials (without startle probes) were used for PCA estimation in the CS-locked window, in order to preserve comparability with the primary analysis.

(2) *Functional out-of-sample validation* was performed without re-estimating the scoring solution, i.e., component scores were projected onto all valid trials based on fixed PCA coefficients obtained from the primary sample. As in the main analysis, trial-wise component scores were analyzed using LMMs assessing design-based a-priori contrasts (with random intercepts for subjects and samples). Due to the overlap of later phases of the CS-locked window with actual or expected US onset in several external studies (see above), results for Post-CS3 scores should be interpreted with caution. Moreover, because US valence was confounded with study characteristics in the external dataset, valence-dependent effects should be evaluated in terms of converging evidence for the temporal pattern identified in the primary analysis (i.e., generalization test) rather than interpreted as a fully independent replication.

**Results.** (1) *Structural replication*: Each four-component PCA solution explained > 80% of total variance in pupillary waveforms (CS-locked window: 87.7%; US-locked: 93.5%). As illustrated in Figure S10, the temporal loading patterns of the reference solutions and the respective PCA of the external dataset corresponded almost perfectly in both windows (Tucker φs ≥ .99). Likewise, indices of both split-half congruence and between-study congruence (based on an iterative leave-one-sample-out analysis) were very high for both windows of analysis (all Tucker φs ≥ .97), confirming excellent stability of the given PCA solution within the external set of studies as well.

(2) *Functional validation*: The pattern of fixed-effects contrasts emerging in LMMs of component scores (see Table S6) largely reproduced the functional dissociations observed in the primary set of studies, including preferential appetitive response differentiation during the early (Post-CS1) phase (with a trend-level valence contrast; see below) and preferential aversive response differentiation around expected outcome onset. Temporal specificity was (very likely) also preserved, as no systematic effects emerged for the Pre-CS component, and *US presence* did not significantly (and clearly not positively) modulate the Pre-US component, whereas pronounced outcome effects emerged in both post-US components. In later CS-locked components, the results deviated somewhat from the original findings, with aversive effects being substantially larger in external studies, which is consistent with the markedly earlier average US onset. The pattern of results was largely consistent across samples (for an overview, see Figure S11).

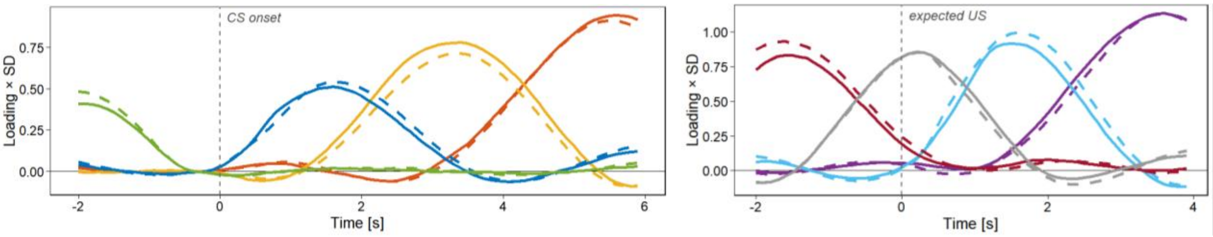


**Supplementary Figure S10.** Loading patterns of PCA solutions based on external data (solid lines) in comparison to the respective reference solution (dashed lines), as described in the main text, in the CS-locked (left) and US-locked (right) windows.

While the valence-dependent contrast was only marginally significant for Post-CS1 scores, pairwise comparisons indicated that the (significant) early CS+ vs. CS- contrast was mainly driven by the difference between appetitive CS+ and CS- ($b$ = 0.069, $SE$ = 0.022, $z$ = 3.18, $p$ = .005 [Bonferroni-Holm corrected]), whereas differentiation for aversive CS+ was numerically much smaller and statistically non-significant ($b$ = 0.021, $SE$ = 0.018, $z$ = 1.18, $p$ = .239). By contrast, differentiation in later CS-locked responses was only significant for aversive learning ($p$s < .0001) but not for appetitive CS+ ($p$s > .43). At the same time, the stronger extent of unconditioned responding previously observed in appetitive conditioning was even significant for early post-US responses (in addition to Post-US2 scores) and overall more pronounced than in the primary dataset (see Table S6 and Figure S11). Thus, the strong outcome sensitivity of both post-US components generalized robustly, whereas the magnitude and temporal pattern of its valence modulation differed substantially from the primary dataset.

**Supplementary Table S6.** Comparison of model parameters (main contrasts) of linear mixed-effects models (for each component) in the primary analysis with data used for external validation

| Component | Contrast (fixed effect) | Primary analysis | External validation | |
|---|---|---|---|---|
| | | *b* | *b* [95% CI] | *p* |
| ***Pre-CS*** | *CS+ vs. CS-* | -0.016 | -0.006 [-0.035, 0.024] | .705 |
| | *App. vs. avs.* \| *CS+* | 0.024 | 0.032 [-0.022, 0.085] | .247 |
| ***Post-CS1*** | *CS+ vs. CS-* | 0.012 | 0.045 [0.017, 0.073] | .001 |
| | *App. vs. avs.* \| *CS+* | 0.078 | 0.048 [-0.007, 0.103] | .083 |
| ***Post-CS2*** | *CS+ vs. CS-* | 0.048 | 0.122 [0.091, 0.153] | < .001 |
| | *App. vs. avs.* \| *CS+* | 0.002 | -0.207 [-0.268, -0.145] | < .001 |
| ***Post-CS3*** | *CS+ vs. CS-* | 0.040 | 0.229 [0.197, 0.260] | < .001 |
| | *App. vs. avs.* \| *CS+* | -0.008 | -0.438 [-0.499, -0.377] | < .001 |
| ***Pre-US*** | *CS+ vs. CS-* | 0.037 | 0.058 [0.031, 0.086] | < .001 |
| | *App. vs. avs.* \| *CS+* | -0.009 | -0.065 [-0.120, -0.011] | .019 |
| | *US+ vs. US-* \| *CS+* | -0.021 | -0.031 [-0.070, 0.008] | .119 |
| | *App. vs. avs.* \| *US+* | -0.024 | 0.025 [-0.053, 0.103] | .527 |
| ***Peri-US*** | *CS+ vs. CS-* | 0.059 | 0.085 [0.055, 0.115] | < .001 |
| | *App. vs. avs.* \| *CS+* | -0.074 | -0.402 [-0.461, -0.343] | < .001 |
| | *US+ vs. US-* \| *CS+* | 0.021 | -0.057 [-0.100, -0.015] | .008 |
| | *App. vs. avs.* \| *US+* | -0.097 | -0.134 [-0.219, -0.050] | .002 |
| ***Post-US1*** | *CS+ vs. CS-* | 0.250 | 0.453 [0.425, 0.480] | < .001 |
| | *App. vs. avs.* \| *CS+* | -0.074 | -0.019 [-0.073, 0.036] | .501 |
| | *US+ vs. US-* \| *CS+* | 0.488 | 0.724 [0.685, 0.763] | < .001 |
| | *App. vs. avs.* \| *US+* | 0.010 | 0.353 [0.275, 0.431] | < .001 |
| ***Post-US2*** | *CS+ vs. CS-* | 0.203 | 0.256 [0.226, 0.287] | < .001 |
| | *App. vs. avs.* \| *CS+* | 0.017 | 0.184 [0.130, 0.239] | < .001 |
| | *US+ vs. US-* \| *CS+* | 0.396 | 0.503 [0.460, 0.546] | < .001 |
| | *App. vs. avs.* \| *US+* | 0.081 | 0.477 [0.391, 0.563] | < .001 |

*Note*. Estimates from the primary analysis (see main text) are reproduced for direct comparison. Estimates for the external dataset were obtained from the prespecified LMMs applied to component scores projected using the respective primary PCA scoring solution (temporal loading pattern). Note that using the external PCA for projection would yield equivalent results. App., appetitive; avs., aversive.

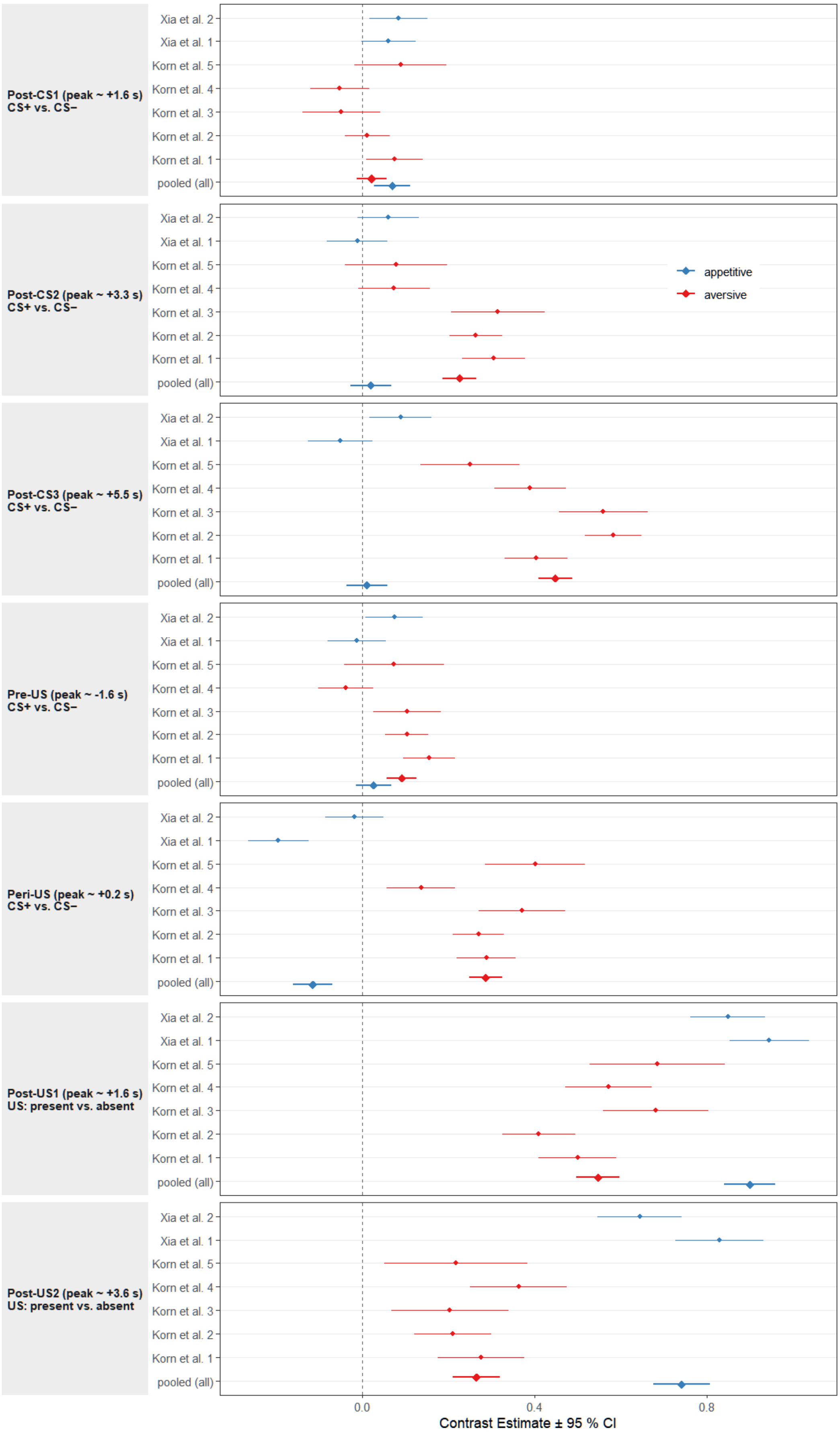


**Supplementary Figure S11.** Distribution of effects (main contrasts) across samples in the dataset used for external validation, shown separately for each component.
